# Competing effects modulate the rate of poly(A) RNA deadenylation in a biomolecular condensate

**DOI:** 10.64898/2026.07.02.736149

**Authors:** Rose M. Irwin, Robert W. Harkness, Brian Tsang, Zi Hao Liu, Kunyang Sun, Tian Hao Huang, Teresa Head-Gordon, Lewis E. Kay, Julie D. Forman-Kay

**Affiliations:** Program in Molecular Medicine, Hospital for Sick Children; Toronto, ON, M5G 0A4, Canada; Department of Biochemistry, University of Toronto; Toronto, ON, M5R 0A3, Canada; Department of Molecular Genetics, University of Toronto; Toronto, ON, M5R 0A3, Canada; Department of Chemistry, University of Toronto; Toronto, ON, M5S 3H6, Canada; Department of Chemistry, Bioengineering, and Chemical and Biomolecular Engineering, University of California, Berkeley; Berkeley, CA, 94720, USA; Chemical Sciences Division, Lawrence Berkeley National Laboratory; Berkeley, CA, 94720, USA

**Author notes:** Department of Molecular and Cellular Biology, University of Guelph; Guelph, ON, N1G 2W1, Canada. Department of Diagnostic Radiology, The Ottawa Hospital – Civic Campus; Ottawa, ON, K1Y 4E9, Canada.

**Keywords:** biomolecular condensate, enzyme, kinetics, phase separation, CNOT7, CAPRIN1, deadenylation, poly(A), RNA

## Abstract

The unique solvent milieu found in biomolecular condensates can control cellular enzymatic reactions and shift reaction kinetics by modulating reactant concentrations, structural dynamics, and enzyme activities. Here we explore the interplay of multiple regulatory factors within a condensate to control poly(A) RNA deadenylation, the first and rate-limiting step in mRNA turnover. The deadenylase CNOT7, a subunit of the CCR4-NOT deadenylation complex, localizes to cytoplasmic RNA granules and shows increased degradation activity *in vitro* in condensates formed by the C-terminal low complexity disordered region of CAPRIN1, a component of RNA granules. We use a combination of enzymatic assays, kinetic modeling, microscopy, Nuclear Magnetic Resonance (NMR) spectroscopy, and molecular dynamics simulations to deconvolute and define the components that underlie this enhancement. We found that enzyme and RNA are concentrated in condensates relative to buffer, which increases CNOT7 activity, while the equilibrium between CNOT7’s active and inactive states remains unchanged. The concentration-dependent increase in enzymatic rates is counterbalanced by a substantial decrease in the enzyme’s catalytic efficiency, likely due to slower diffusion of CNOT7 and RNA within the condensates, which lessens the probability of enzyme-substrate complex formation. Molecular dynamics simulations reveal CNOT7-CAPRIN1 interactions that rely on conserved CAPRIN1 sequence features, hinting at an evolutionarily conserved role for CAPRIN1 condensation. With this quantitative kinetic analysis, we describe the multifaceted mechanism behind regulation of CNOT7 deadenylation by a condensate environment.

**GRAPHICAL ABSTRACT:** 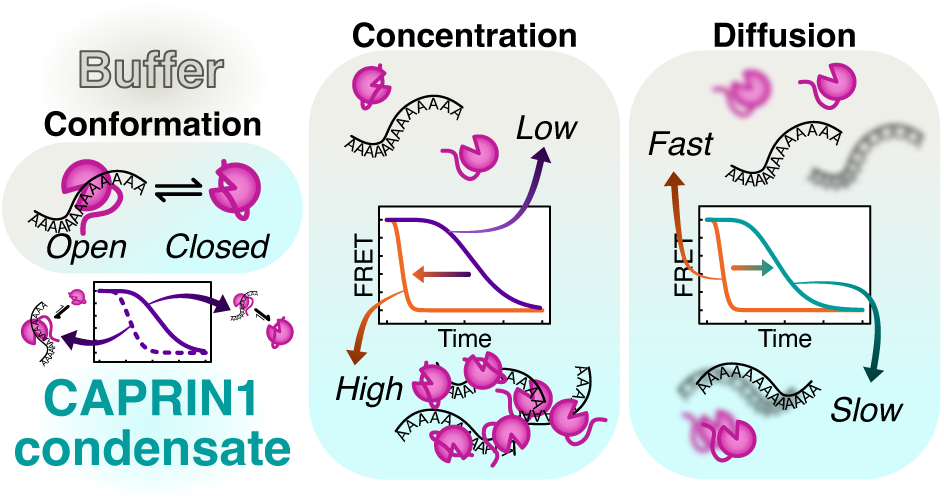

## INTRODUCTION

Biomolecular condensates are increasingly recognized for their roles in controlling cellular enzymatic reactions^1–4^. These include DNA repair by DNA damage foci^5,6^, RNA degradation by processing bodies (P-bodies) and bacterial ribonucleoprotein (RNP) bodies^7,8^, and translation by stress granules and RNA transport granules^8,9^. These regulatory roles are facilitated by the varying physicochemical, viscoelastic, and network properties that emerge within biomolecular condensates. Condensates can form through intermolecular interactions between numerous types of molecules, such as proteins, nucleic acids, metabolites, and ions^10^. Condensate assembly leads to molecular partitioning (enrichment or exclusion) depending on the species involved and can occur with marked changes in the concentrations of the molecules dissolved within them^11–14^ . The unique properties within condensates, including interaction-network percolation, increased crowding, and changes to solvent and material properties such as pH, polarity, viscosity, elasticity, and permeability, for example^12,13,15–20^, impact the stability and activity of the condensate-solvated biomolecules^21–26^.

Compartmentalization via biomolecular condensates can regulate enzymatic activity and substrate selectivity by multiple mechanisms, one example of which is simply modulating concentrations^13,27–29^. Enzymatic reaction rates can increase by concentrating substrates with the enzyme^29–31^ or increasing enzyme cofactor recycling^32^. Partitioning also drives downregulation of activity in cases where substrates are excluded from the condensate^29,33^. Decreased activity has been correlated to reduced diffusion in condensates^34^. In another case, phase separation driven by network formation via homotypic interactions favors the inactive state of an enzyme. This inhibition is alleviated by the addition of other proteins that alter the interaction network and reduce the population of inactive enzyme^35^.

The physicochemical environment within a biomolecular condensate can also impact enzyme kinetics by influencing substrate-enzyme binding affinities or conformational equilibria essential for enzymatic activation^31,35,36^. Binding affinity between enzyme and substrate can potentially be enhanced depending on the network formed by scaffold proteins within the condensate^31^. Enzymes may also display increased activity due to changes in thermodynamic stability^36^.

A number of studies have broadly quantified the effects of condensates on enzymatic activity, primarily using engineered systems to form condensates and/or localize the enzyme into the dense phase^31,34,36^. Here we sought to quantify how the solvent environment of a biomolecular condensate would affect the activity of an enzyme that inherently (*i.e.*, without the addition of a dense-phase localization tag^13,34^) partitions into it. Therefore, we exploited an *in vitro* model system previously studied by our lab in exploration of poly(A) RNA deadenylation kinetics^16,33,37–39^. The condensate is formed by the C-terminal low-complexity region (LCR) residing in an intrinsically disordered region (IDR) of the RNA-binding protein and translational regulator Cytoplasmic Activation- and Proliferation-Associated Protein 1 (CAPRIN1 607-709, referred to here as CAPRIN1). Full length CAPRIN1 is highly enriched in the brain^40,41^ and localizes to cytoplasmic RNA granules including P-bodies, stress granules, and mRNA transport granules^42–44^. The shortened construct provides a simple model for these RNA regulatory condensates.

The enzyme-substrate system within CAPRIN1 condensates used here is comprised of the mRNA deadenylase CNOT7, a vital subunit of the eukaryotic deadenylase CCR4-NOT complex^45,46^, and poly(A) RNA. CNOT7 is essential for cell viability^47,48^ and influences synaptic plasticity in cultured neurons by regulating dendritic mRNA transport and local translation^49^. As a deadenylase, CNOT7 reduces poly(A) tail length, which dictates mRNA turnover and translational efficiency^50^, and its deadenylation activity may be regulated by stress granule assembly^51^. As such, understanding the influence of a condensate environment on CNOT7 activity will aid in developing a detailed picture of mRNA homeostasis and regulation of gene expression.

We previously found that, for the same bulk concentrations of enzyme and substrate, the rate of deadenylation in phase-separated CAPRIN1 conditions was faster than in buffer, although an explanation for this enhancement was not pursued^33^. Here we elucidate mechanisms underlying the observed enhancement through enzymatic assays, reaction rate modeling, Nuclear Magnetic Resonance (NMR) spectroscopy, and molecular dynamics simulations. Our analyses evaluate three potential factors regulating deadenylation rates that can combine to yield the observed rate acceleration in CAPRIN1 condensates under the conditions we have explored. These include dynamics of the C-terminal tail of CNOT7 controlling accessibility of the enzyme’s active site to RNA, increased dense phase enzyme and substrate concentrations, and modulation of CNOT7’s catalytic efficiency. Although our focus is on the specific case of deadenylation, the combination of these effects underscores the complexity and versatility of biomolecular condensates as solvents that regulate cellular processes.

## RESULTS

### Modulation of CNOT7 activity via C-terminal tail dynamics

To better understand deadenylation by CNOT7, we sought to clarify potential regulatory mechanisms of the enzyme. In one of the crystal structures of CNOT7 (PDB 4GMJ)^52^, the C-terminal tail occupies the RNA-binding site (Figure 1A). The tail contains three consecutive glutamates (278-280) that, in this structure, coordinate the active site magnesium ions, which are necessary for RNA degradation activity^53^. The tail also contains two tyrosine residues that, along with the glutamates, could participate in electrostatic, π-π, and/or cation-π interactions with the positively charged (+12.8 at pH 7.4) arginine-/aromatic-rich CAPRIN1 or RNA. We therefore hypothesized that the tail influences deadenylation activity by modulating access of RNA to the active site, with interactions with CAPRIN1 in the condensate potentially biasing the tail conformations to enhance RNA binding.

**Figure 1.**
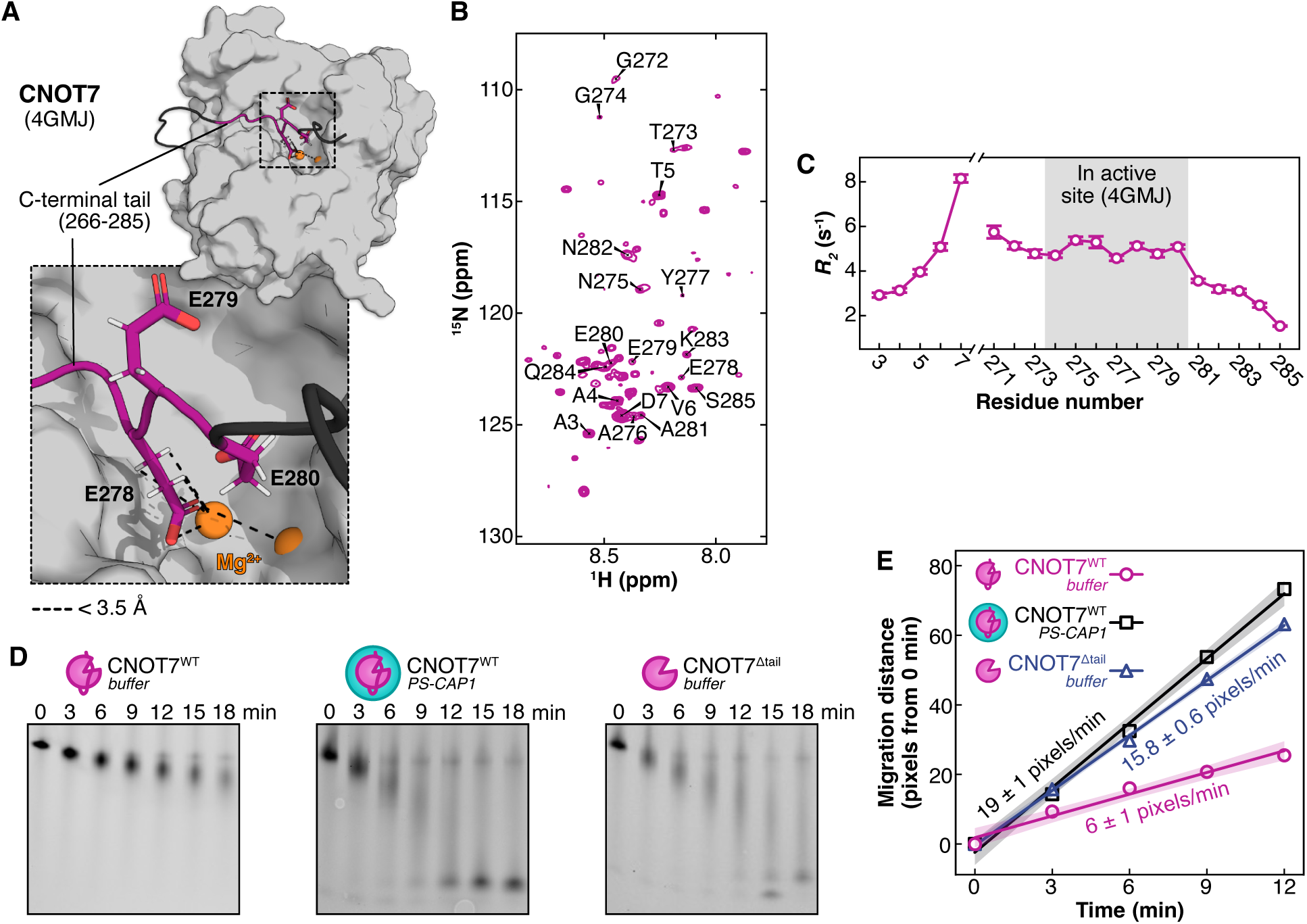
CNOT7 activity is modulated by its C-terminal tail and partitioning into CAPRIN1 condensates. (A) CNOT7 crystal structure (PDB 4GMJ chain B) with the folded domain in grey, resolved C-terminal tail in pink, and missing tail residues built by IDPConformerGenerator^54^ in black. Dashed lines indicate distances of ≤ 3.5 Å between magnesium ions (orange) and tail residues in the active site (inside the black square). (B) ^15^N-^1^H HSQC spectrum of CNOT7^WT^ with N- and C-terminal tail assignments, 30 °C, 800 MHz. (C) Transverse relaxation rates *R_2_* (s^-1^) for CNOT7^WT^ with 2 mM Mg^2+^, 30 °C. The grey region indicates tail residues resolved in the 4GMJ crystal structure. Error bars indicate one standard deviation from the mean of two experiments in measured *R_2_* rates. (D) RNA degradation by CNOT7^WT^ at 30 °C in buffer (*left*) and within condensates (PS-CAP1, *middle*), and by CNOT7^Δtail^ in buffer (*right*). Bulk concentrations were 1 µM enzyme, 0.3 µM poly(A) RNA(A)_38_, and 100 µM CAPRIN1 when present (see Figure S1A for gels with RNA(A)_38_ concentrations up to 5 µM). (E) Migration distances from gels in D (Δpixels) as a function of time, with linear fits. The slope of each fit (reported on the plot ± standard error of the fit) approximates the deadenylation rate in pixels/min. The shaded areas surrounding each fitted line correspond to the 95% confidence intervals.

We first employed NMR spectroscopy to study the C-terminal tail in solution, which is expected to be dynamic relative to the folded region of CNOT7. The ^15^N-^1^H HSQC spectrum of CNOT7 (Figure 1B) has sharp resonances for the N- and C-terminal tail residues, with a narrow distribution of amide proton positions, indicating disorder. In contrast, resonances for the folded catalytic domain are weaker, consistent with overall slow tumbling for the well-folded region of the protein and likely conformational exchange within the folded domain. We found that the transverse relaxation rates (*R_2_*), which are proportional to the NMR resonance linewidths, for both the N- and C-termini were relatively low and significantly lower than for residue 7, the last residue for which a sharp amide resonance was observed before the start of the folded domain (Figure 1C). This suggests that the tail samples a dynamic ensemble of conformations that are not stably interacting with the active site.

To test the impact of the tail on deadenylase activity in a condensed phase, we performed preliminary experiments using a gel-based assay^55^ at 30 °C with a synthetic poly(A) RNA substrate fluorescently tagged with fluorescein isothiocyanate (FITC), 5’ FITC-CCUUUCC(A)_38_, denoted as RNA(A)_38_ (Figures 1D, and S1A). In what follows, we refer to the one-phase, CAPRIN1-free condition as “buffer” and conditions with phase-separated CAPRIN1 as “PS-CAP1”. The initial rates of reaction were quantified with FITC signal migration over time for wild-type CNOT7 (CNOT7^WT^) in buffer, or with phase-separated CAPRIN1, as well as a truncated CNOT7 construct in which the C-terminal tail was removed, CNOT7^Δtail^, in buffer (Figure S2).

At 0.3 µM RNA(A)_38_, the apparent rates of reaction for 1 µM CNOT7^Δtail^ in buffer conditions and 1 µM CNOT7^WT^ in PS-CAP1 conditions were similar and ∼3-fold faster than for CNOT7^WT^ in buffer (Figure 1E, see Figure S1B for quantification of all conditions). This trend is observed below ∼2 µM of RNA(A)_38_ for which there is significant deadenylation over the short time course of the reaction (15 minutes) (Figure S1C). These findings are consistent with a previously proposed picture where the CNOT7^WT^ tail samples conformations near the active site in buffer, resulting in a net attenuation of adenylation, likely by restricting RNA binding^56^. These data could suggest that under PS-CAP1 conditions the tail dynamics are biased toward an ensemble that does not impede the RNA from the enzyme active site. However, there are multiple factors that must be considered when the reaction is performed in condensates, including local concentrations of enzyme and substrate relative to buffer, as well as the catalytic efficiency of the enzyme in the dense proteinaceous droplet environment.

### CNOT7 conformational equilibrium and rate parameters in buffer conditions

Our gel-based assays establish that the CNOT7 tail decreases the rate of deadenylation relative to a tailless enzyme (Figures 1E and S1B, compare CNOT7^WT^ and CNOT7^Δtail^ in buffer conditions at 0.3 µM RNA(A)_38_ where an approximate 3-fold difference is observed). The presence of the tail in the active site in one of the X-ray structures of the enzyme, occluding RNA from binding, together with our NMR data demonstrating a disordered tail, suggested that CNOT7 might exchange between ‘inactive’ (tail bound to RNA-binding site) and ‘active’ (tail free from the RNA-binding site) conformations in solution to modulate activity. To explore this, we conducted detailed kinetic assays to measure cleavage rates of poly(A) RNA using a Förster Resonance Energy Transfer (FRET)-based assay described in detail previously^57^, focusing initially on deadenylation in the buffer reference condition to establish a baseline for further analyses of the effects of CAPRIN1 on the reaction.

Figure 2 illustrates the experimental workflow that was used. Briefly, aliquots of enzyme and RNA were distributed in a 384-well plate reader and the reaction allowed to proceed for a given period before quenching. Notably, the quenching buffer also included carboxytetramethylrhodamine (TAMRA)-labeled poly(T) DNA (5’ (T)_38_GGAAAGG-TMR) that annealed with the RNA products in a manner dependent on the length of the RNA. The FRET between the TAMRA (abbreviated in the figure as TMR, on the DNA) and 6FAM (on the RNA) fluorescence labels provides a readout of the reaction progress, with the signal decreasing as the RNA is degraded because the stability of the DNA-RNA hybrid is reduced (see *Supporting Information: ‘FRET-based Deadenylation Assay’*).

**Figure 2.**
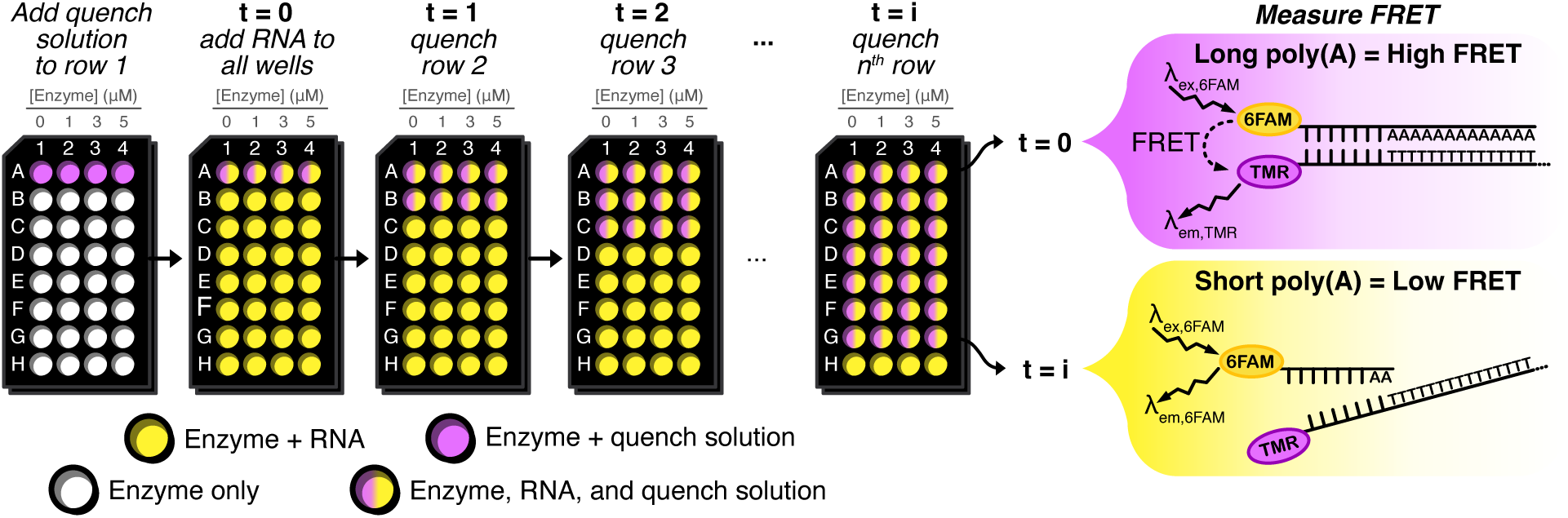
Schematic of the FRET-based deadenylation assay protocol. Enzyme was mixed with RNA(A)_18_ (yellow), a poly(A) RNA substrate that has a 5’ conjugated 6FAM fluorophore and a poly(A) tail of 18 adenosines (5’ 6FAM-CCUUUCC(A)_18_). The reaction was quenched at regular time points by adding 1% SDS, 0.1 mg/mL Proteinase K, and a 3’ TAMRA fluorophore-conjugated DNA strand (5’ (T)_38_GGAAAGG-TMR, DNA(T)_38_) (pink) that is complementary to the full-length RNA substrate and the deadenylation reaction intermediates. If the RNA is long, the RNA and DNA anneal with high affinity and produce high FRET – *i.e.*, high signal from the DNA(T)_38_ (pink). If the RNA has been degraded, annealing is diminished, higher 6FAM fluorescence (yellow) is observed, and the FRET decreases (see *Supporting Information: ‘FRET-based Deadenylation Assay’*).

To describe these FRET-based datasets, we developed an enzymatic model which includes a CNOT7^WT^ inactive-active pre-equilibrium to fit the resulting activity profiles (Scheme 1, see *Supporting Information: ‘Deadenylation kinetic model’*).

**Scheme 1.** Enzymatic model used to describe FRET assay data.

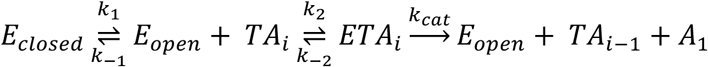

Parameters *k*_1_ and *k*_–1_ are rates for the release and binding of the C-terminal tail, respectively, to form the active and inactive enzyme (*E_open_* and *E_closed_*, respectively). The rates of association of the active (open) enzyme with poly(A) tails of length *i* (*TA_i_*) and dissociation of enzyme-RNA complexes (*ETA_i_*) are given by *k_2_* and *k_-2_*, respectively. The rate of the catalytic step, *k_cat_*, quantifies removal of a single adenosine *A*_1_ from an enzyme-bound RNA to generate free enzyme and a shorter RNA product *TA_i_*_–1_. Note that *T* is a tag that positions the annealed RNA and DNA strands appropriately so that the attached fluorophores to the two strands are aligned appropriately for FRET (see Figure 2 and Supporting Information).

In this model, CNOT7 exchanges between closed, inactive (*E_closed_*, tail bound) and open, active (*E_open_*, tail unbound) conformations, with only the open state, *E_open_*, able to bind the RNA substrate leading to deadenylation. Fits of simulated data (Figure 3A) show that the output individual model parameters are poorly defined and correlated with their starting values (Figure 3B). In contrast, a parameter defined as the observed reaction rate, *k*^∗^, with

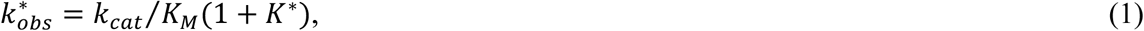

**Figure 3.**
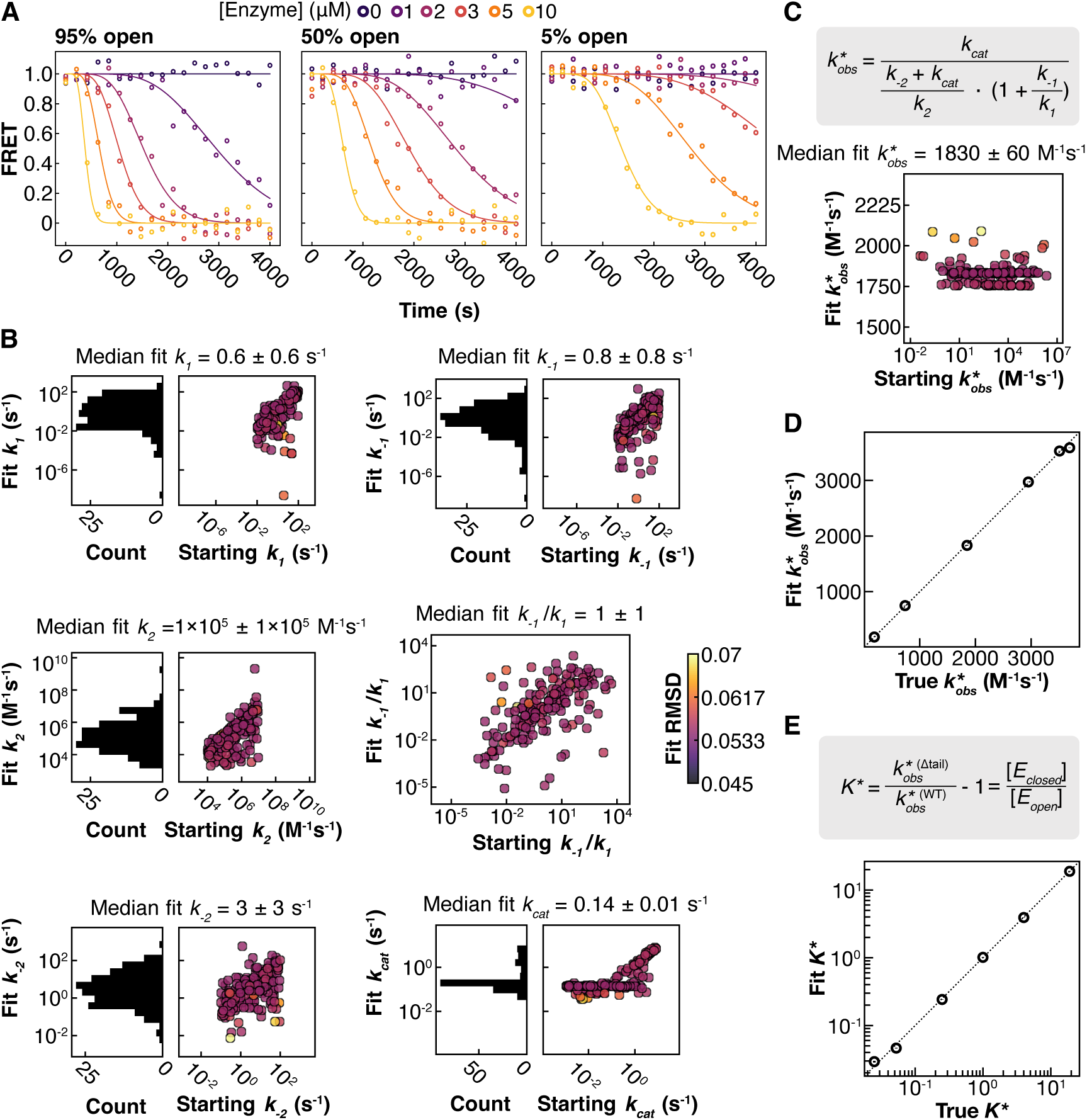
Simulations show that accurate values of 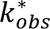 and *K*^∗^ are obtained from analysis of the FRET assays. (A) Simulated deadenylation assay data with 100 nM RNA(A)_18_ and 0-10 µM enzyme with the rate constants *k*_1_, *k*_2_, *k*_–2_, and *k_cat_* (Scheme 1) set to 1 s^-1^, 1×10^5^ M^-1^ s^-1^, 3 s^-1^, and 1.15×10^-1^ s^-1^, respectively, while *k*_–1_ = 5.26×10^-2^ s^-1^, 1 s^-1^, and 19 s^-1^ to generate an array of different fractions of open CNOT7 at equilibrium (95%, 50%, and 5%, respectively). The rate values were chosen because they produce FRET curves like those measured experimentally. Noise was added to the data based on the average RMSD of all experimental fits (0.056). Simulated data were fit with 200 sets of starting parameters randomly distributed across several orders of magnitude (*k*_1_ = 10^-2^-10^2^ s^-1^, *k*_–1_ = 10^-2^-10^2^ s^-1^, *k*_2_ = 10^4^-10^7^ M^-1^s^-1^, *k*_–2_ = 10^-1^-10^2^ s^-1^, and *k_cat_* = 10^-3^-10^1^ s^-1^). Fits with an RMSD < 7×10^-2^ were used for further analysis. (B) Example starting fit (x-axis) and final fit (y-axis) rate values from simulations with *K*^∗^ = 1.0 (50% open); 179 out of 200 fits achieved an RMSD ≤ 7×10^-2^ (see A for visual). Data points in the correlation plots of final fit *vs* starting fit parameter values are colored according to the RMSD of the fit (see scale bar on the right of B). The histogram to the left of each plot shows the final fit values on the same scale as the correlation plot y-axis. Median final fit values ± median absolute deviation are reported above each plot, demonstrating that individual parameters are poorly defined. (C) Starting fit and output fitted 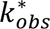 (M^-1^s^-1^) rates were calculated for each fit (equation shown in grey box in C, see eq 1). Although the starting fit input 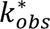 values spanned several orders of magnitude, the resultant fitted values were nevertheless in good agreement with each other and with the ground truth 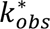 value used to generate the profiles (1.82×10^3^ M^-1^s^-1^). The median fitted 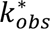 ± median absolute deviation is reported above the plot. (D) The analysis shown in B was repeated for data simulated with 5%, 20%, 80%, 95%, and 100% open enzyme. Median fit 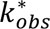 values were then determined from the values for all fits with an RMSD ≤ 7×10^-2^ and plotted against ground truth 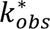 values. Error bars indicate median absolute deviation. The dashed line y = x is shown to guide the eye. (E) Median fit-derived *K*^∗^ values (equation shown in grey box) for all simulated datasets plotted against their ground truth *K*^∗^ values, calculated by taking the ratios of 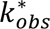 values obtained from fits of simulations of FRET data for CNOT7^WT^ and CNOT7^Δtail^. Note that for CNOT7^Δtail^ a value of *K*^∗^ = 0 is used, as the absence of a tail is equivalent to the ‘all out’ tail conformation. Error bars indicate median absolute deviation. The dashed line y = x is plotted.

is obtained directly from our analysis (see *Supporting Information: ‘Deadenylation kinetic model’*) and is well defined (Figure 3C and 3D), and from it, robust measures of the open/closed equilibrium, *K*^∗^ = [*E_closed_*]/[*E_open_*], can be extracted (Figure 3E; see *Supporting Information: ‘Determining the accuracy of the rate constant k*^∗^ *and equilibrium constant K*^∗^*’*). In eq 1, the Michaelis constant, *K_M_*, is given by *K_M_* = (*k*_–2_ + *k_cat_*)⁄*k*_2_. Furthermore, simulations confirm that *ETA_i_* (from Scheme 1) is effectively in steady state because our experiments are performed with enzyme concentrations greatly exceeding the (fixed) substrate concentration, and that the concentrations of *E_closed_* and *E_open_* are constant over the reaction trajectory (Figure S3; see *Supporting Information: ‘Evaluating the steady state assumption and assessing the ratio of E_closed_ and E_open_’*). Unlike measured rate *vs* time profiles, which depend on concentrations of enzyme and substrate, the fitted 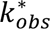 values are independent of concentrations of reactants and products, enabling changes in equilibria associated with tail in/out conformations and changes due to catalytic efficiency to be assessed.

Using the RNA substrate RNA(A)_18_ (5’ 6FAM-CCUUUCC(A)_18_), having a 6-carboxyfluorescein (6FAM) label, we measured rate profiles for CNOT7^WT^ and CNOT7^Δtail^ in buffer conditions at 25 °C (Figures 4A and S4) and determined 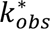 values (Figures 4B and S5, Table S1) for each construct. The 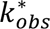 value for CNOT7^Δtail^ in buffer was higher than that of CNOT7^WT^ in buffer (3300 ± 500 s^-1^ *vs* 1900 ± 200 s^-1^), consistent with results from the more qualitative gel-based assay where CNOT7^Δtail^ was faster than CNOT7^WT^ (in conditions with [RNA(A)_38_] < 2 µM; Figures 1E and S1B). By taking CNOT7^Δtail^ as the fully open state (*K*^∗^ = 0) and assuming that *k_cat_*⁄*K_M_* does not vary for the two CNOT7 constructs for a given set of experimental conditions (*i.e.*, same temperature and comparing results from both CNOT7 variants in either buffer or in PS-CAP1, for example), it is possible to calculate *K*^∗^ for CNOT7^WT^ from the ratio of 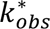 values for CNOT7^WT^ and CNOT7^Δtail^ (Figure 3E, eq S26). In this manner a value of *K*^∗^ = 0.7 ± 0.2 was calculated for CNOT7^WT^ (Figures 4C and S7, Table S1; see *Supporting Information: ‘Deadenylation kinetic model’*) at 25 °C, corresponding to a fractional population of 42 ± 5% in the closed conformation (Figure 4D). These results underscore the importance of the disordered tail in regulating CNOT7 activity.

**Figure 4.**
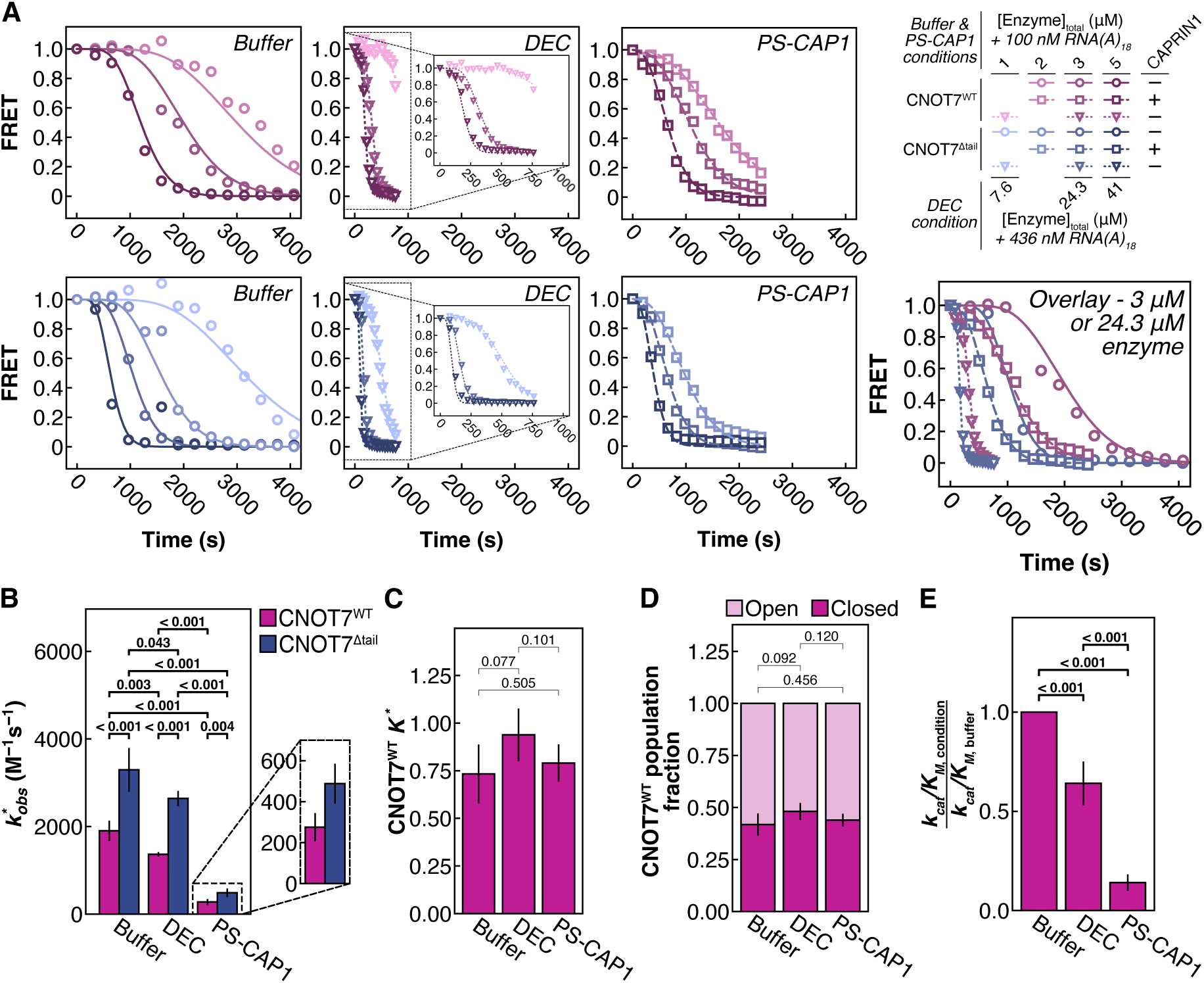
CNOT7^WT^ activity in CAPRIN1 condensates is modulated by increases in enzyme and RNA concentrations and reduced catalytic efficiency. (A) Representative data and fits for degradation of 6FAM-RNA(A)_18_ by CNOT7^WT^ (pink) and CNOT7^Δtail^ (blue) in buffer (circles, solid lines), droplet-equivalent concentration (DEC, inverted triangles, dotted lines), or in phase-separated CAPRIN1 (100 µM bulk CAPRIN1, PS-CAP1, squares, dashed lines) conditions at 25 °C. Deadenylation rates for buffer conditions were based on enzyme concentrations ranging from 1-7 µM (1-5 µM shown here; see Figure S4) and 100 nM RNA. DEC activity was measured in the absence of CAPRIN1 at enzyme concentrations corresponding to those in the dense phase of PS-CAP1 samples prepared with 1, 3, and 5 µM bulk enzyme concentrations, corresponding to 7.6 µM, 24.3 µM, and 41 µM enzyme within the condensates, respectively. Due to minimal difference in dense-phase RNA(A)_18_ concentrations as a function of enzyme (Table S3), the RNA concentration for DEC assays was set to 436 nM, the average dense-phase RNA concentration across measured conditions. PS-CAP1 data with 2-5 µM total enzyme and 100 nM total RNA(A)_18_ were fit with a two-phase model to isolate the 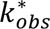 in the dense phase (see *Supporting Information: ‘Fitting PS-CAP1 deadenylation assay data with a two-phase model’*). See Figure S4 for replicates. All data were background corrected based on a 0 µM enzyme condition. Overlay shows the data and fits for 3 µM bulk enzyme/100 nM bulk RNA(A)_18_ (buffer, PS-CAP1) or 24.3 µM enzyme/436 nM RNA(A)_18_ (DEC) for comparison. (B) Average rate of deadenylation 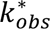 (M^-1^s^-1^) for CNOT7^WT^ (pink) and CNOT7^Δtail^ (blue) in buffer, DEC, or PS-CAP1 conditions for data in A and Figure S4 (see Figure S5 for individual replicate 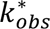 values). Note that although the extracted 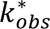 for buffer and DEC are similar, the DEC profiles in A decay more rapidly owing to the elevated enzyme concentrations. (C) Average CNOT7^WT^ conformational equilibrium *K*^∗^ in buffer, PS-CAP1, or DEC conditions (see Figure S5 for individual replicate *K*^∗^ values). (D) Fraction of CNOT7^WT^ open (light pink) or closed (dark pink) for each condition based on *K*^∗^ values in C and eq S27. (E) The ratio between CNOT7 *k_cat_*/*K_M_* (catalytic efficiency, calculated from 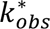 values obtained from measurements using the CNOT7^Δtail^ variant where *K*^∗^ = 0) in the indicated condition (x-axis) normalized to *k_cat_*/*K_M_* in buffer (eq S32). For all panels, error bars show one standard deviation from the mean, and *P* values for unpaired *t* tests are shown above bars with significant values (*P* < 0.05) bolded.

### Increase in CNOT7 activity due to high concentration in CAPRIN1 condensates

Having measured the deadenylation reaction for both CNOT7^WT^ and CNOT7^Δtail^ and quantified the in/out tail equilibrium, *K*^∗^, for CNOT7^WT^ in buffer, we next proceeded to measure deadenylase activities of CNOT7 in PS-CAP1 conditions (100 µM bulk CAPRIN1 concentration) using the FRET-based assay. We first confirmed that CNOT7^WT^ and CNOT7^Δtail^ partition into condensates with RNA(A)_18_ and CAPRIN1 via confocal microscopy (Figure S6).

We then quantified dense- and dilute-phase concentrations of enzyme and substrate to enable a direct comparison of reaction rates in buffer and PS-CAP1 conditions, as has been done in previous studies^31,34^. For these measurements, we imaged phase-separated condensates comprised of CNOT7^WT^ or CNOT7^Δtail^ with 1, 3, or 5 µM total enzyme (with 50 nM Alexa Fluor 647-enzyme) and 100 nM 6FAM-RNA (Figure S7A) and determined the total, dense-phase, and dilute-phase concentrations of labeled enzyme and RNA. For a given total enzyme concentration, we observed no clear differences between CNOT7^WT^ and CNOT7^Δtail^ concentrations in the dense phase, and similarly in the dilute phase, and the concentrations of RNA in the dense and dilute phases were also found to be the same when either CNOT^WT^ or CNOT7^Δtail^ was present, so we combined the CNOT7^WT^ and CNOT7^Δtail^ datasets (Figure S7B). Using the average dense phase, dilute phase, and total concentrations of fluorescently labeled enzyme or RNA(A)_18_, we calculated the partition coefficient (PC_labeled_; eq S35) and droplet volume fraction (DVF_labeled_, *i.e*., the fraction of total sample volume made up by the dense phase; eq S36) values for the fluorescently labeled molecules (Figure S7C). We then linearly fit these values as a function of bulk concentration of CNOT7 (1, 3, 5 µM) to interpolate the DVF_labeled_ and PC_labeled_ values for every bulk concentration of enzyme in our PS-CAP1 FRET assays (1, 2, 3, 5 µM; Table S2). Although fluorophores may impact phase behavior^58,59^ (see *Supporting Information: ‘Confocal Microscopy – Limitations’*), droplet volume fraction and partition coefficient values were assumed to be unaffected by the presence of the fluorescent tags in our analyses, and the DVF and PC in assay conditions (*i.e.*, without fluorescently labeled enzyme), DVF_assay_ and PC_assay_, were assumed to be equivalent to DVF_labeled_ and PC_labeled_, respectively. The DVF_labeled_, and the PC_labeled_ for both enzyme and substrate obtained via interpolation were therefore used to calculate dense- and dilute-phase concentrations of both CNOT7 and RNA under the deadenylation reaction conditions of our FRET assays according to eqs S37 and S38 (Table S3).

We measured CNOT7 and RNA(A)_18_ activity at these dense-phase concentrations in a single-phase solution (*i.e.*, without CAPRIN1) to isolate the impact of increased enzyme/substrate concentration on reaction kinetics. The deadenylation assay was run with the droplet-equivalent concentrations (DEC) of enzyme and substrate components as quantified with microscopy in the PS-CAP1 samples (Table S3). We observed significantly faster deadenylation (assessed by time to reach 0 FRET signal, *t_F_*_45_) with the increased amounts of reactants relative to buffer (*t_F_*_45_ ∼ 2300 ± 200 s and 330 ± 50 s, respectively, for 5 µM CNOT7^WT^ in buffer *vs* the equivalent DEC condition, *P* < 0.001; Figures 4A and S4). We then fit our enzymatic model to the DEC data and determined that the 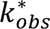 values for CNOT7^WT^ and CNOT7^Δtail^ under DEC conditions were not significantly different than their lower concentration (buffer condition) counterparts (Figures 4B and S5, Table S1). In turn, the calculated *K*^∗^ values for CNOT7^WT^ in buffer *vs* DEC conditions were similar (Figures 4C and S5, Table S1), as were the populations of open/closed enzyme (Figure 4D). Thus, the decrease in *t_F_*_45_ for DEC conditions can be attributed to the increased enzyme and RNA concentrations.

### Lower catalytic efficiency in phase-separated CAPRIN1 conditions

Next, we sought to determine 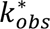 and *K*^∗^ for CNOT7^WT^ in PS-CAP1 conditions. A similar titration of CNOT7^WT^ and CNOT7^Δtail^ (2-5 µM total enzyme with 100 nM RNA(A)_18_) was performed in the presence of 100 µM CAPRIN1 (Figures 4A and S4). The time required to degrade RNA in PS-CAP1 conditions was shorter than under buffer conditions (*t_F_*_45_ ∼ 1800 ± 200 s *vs* 2300 ± 200 s, respectively, for 5 µM total CNOT7^WT^ in PS-CAP1 *vs* buffer conditions, *P* = 0.04) as expected based on previous work^33^ and the gel-based assay, where the rates were faster in PS-CAP1 conditions for [RNA(A)_38_] < 2 µM and 1 µM CNOT7^WT^ (Figure S1C). However, unexpectedly, the time required to degrade the RNA under PS-CAP1 conditions was longer than under DEC conditions (Figures 4A and S4; *t_F_*_45_ ∼ 1800 ± 200 s *vs* 330 ± 50 s, respectively, for 5 µM total CNOT7^WT^ in PS-CAP1 *vs* the equivalent DEC condition, *P* = 0.002), indicating that the condensate environment has an inhibitory influence on activity that counteracts the rate acceleration from elevated enzyme and RNA concentrations.

To understand the origin of the differences in *t_F_*_45_ values, we fit the PS-CAP1 FRET assay data with a two-phase model where deadenylation occurs in both condensed and dilute regions of the sample simultaneously with different kinetic parameters (see *Supporting Information: ‘Fitting PS-CAP1 deadenylation assay data with a two-phase model’*). The two-phase model assumes that (1) a single RNA partition coefficient applies for each RNA species, and (2) all RNA species generated during deadenylation diffuse rapidly between phases to reestablish equilibrium concentrations governed by the RNA DVF and PC values. When solving the RNA degradation rate equations, the total amount of each RNA species was determined after each timepoint and “redistributed” between phases according to eqs S37 and S38. These new concentrations were then used to calculate the further change in RNA species concentrations over the next timepoint, with enzyme concentrations for each phase held constant throughout the fit. During this procedure, the 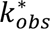 of the dilute phase was fixed to the value measured in buffer, leaving *k*^∗^ for the dense phase as the only adjustable parameter.

Notably, the dense-phase 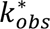values for CNOT7^Δtail^ and CNOT7^WT^ in PS-CAP1 conditions were significantly lower than their corresponding *k*^∗^ values in buffer or DEC conditions (∼7-fold for PS-CAP1 *vs* buffer and 5-fold for PS-CAP1 *vs* DEC; Figures 4B and S5, Table S1). Although we initially hypothesized that in the condensate environment the C-terminal tail of CNOT7 might occupy the RNA-binding site to a lesser extent than in buffer, the extracted *K*^∗^ values for CNOT7^WT^ were similar for all conditions tested (Figures 4C and S5, Table S1), leading to a constant fraction of open and closed enzyme populations (Figure 4D). Therefore, changes in rate cannot be explained in terms of a shift in the tail conformational equilibrium.

What then dictates the observed changes in 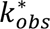? To address this question, we note that 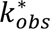 is a function of both *K*^∗^ and the catalytic efficiency, *k_cat_*⁄*K_M_*, and that *k_cat_*⁄*K_M_* can be obtained directly from eq 1 so long as *K*^∗^ is known from a comparison of fitted *k*^∗^ values based on FRET assays recorded using CNOT7^WT^ and CNOT7^Δtail^ (see above and *Supporting Information: ‘Deadenylation kinetic model’*). Values of *k_cat_*⁄*K_M_* for CNOT7^WT^ in both the DEC condition and PS-CAP1 dense phase decreased compared the buffer condition (Figure 4E, Table S4). In particular, the *k_cat_*⁄*K_M_* for CNOT7^WT^ in the PS-CAP1 dense phase decreased to 0.14 ± 0.04 of the *k_cat_*⁄*K_M_* in buffer conditions and 0.2 ± 0.1 of the *k_cat_*⁄*K_M_* in DEC conditions (see *Supporting Information: ‘Evaluating changes in catalytic efficiency’*). Therefore, the increase in overall deadenylase activity based on a comparison of FRET data recorded for the PS-CAP1 and buffer conditions derives from a significant increase in reaction rates due to the higher concentration of enzyme and RNA in the condensate (by ∼7-fold) balanced by a roughly compensatory decrease in catalytic efficiency. In summary, our analysis has enabled an understanding of the impact of the condensate environment on the overall progression of deadenylation, by quantifying the effects of (i) enhancing the concentrations of CNOT7 and RNA, (ii) decreasing the catalytic efficiency, and (iii) monitoring potential changes to the active/inactive CNOT7 conformational equilibrium (Figure 4).

### Solvating interactions between CNOT7 and CAPRIN1

Previous studies have described the distinct solvent properties of CAPRIN1 condensates, specifically increased ion content, decreased concentration of water, and an impact on client protein and RNA stabilities^11,22,26,60,61^. To better understand the physicochemical effects of the CAPRIN1 condensate environment on CNOT7, we performed molecular dynamics simulations (i) in the absence of CAPRIN1 (buffer conditions) and (ii) surrounded by 7 CAPRIN1 LCR chains (residues 607-709; PS-CAP1) to approximate a condensate. For the PS-CAP1 condition, a compaction of CAPRIN1 chains was observed over the 1 µs simulation with the concentration of CAPRIN1 atoms proximal to CNOT7 (in a small box with edge length = 72.7 Å centered around CNOT7, within the overall simulation box with edge length 205 Å) increasing from 10 mM at the start of the trajectory to ∼30 mM (Figure S8A and S8B). At the same time, the concentration of water molecules decreased (Figure S8A), which is consistent with previous experimental work^22,26^ and demonstrates the distinct proteinaceous solvent nature of the simulated CAPRIN1 condensate. A first set of simulations was performed starting from an open CNOT7^WT^ pose, representing the active conformation of the enzyme, with the tail modeled to point away from the active site. Over the course of the simulation in buffer, the tail moved to occupy the active site and remained associated with it subsequently (Figures 5A and S8C). Interestingly, this association was between the Mg^2+^ ion and the terminal carboxyl of S285 (Figure 5A), which suggests multiple modes of tail-active site interaction beyond the EEE-Mg^2+^ contacts observed in the PDB 4GMJ crystal structure (Figure 1A). In contrast, when simulations were performed to mimic the PS-CAP1 environment, the tail resided outside of the active site over the course of the trajectory (Figures 5B and S8C). In another set of simulations performed with a starting pose based on the crystal structure (Figure 1A), minimal tail movement was observed for computations in both buffer and PS-CAP1 solvents (Figure S8C).

**Figure 5.**
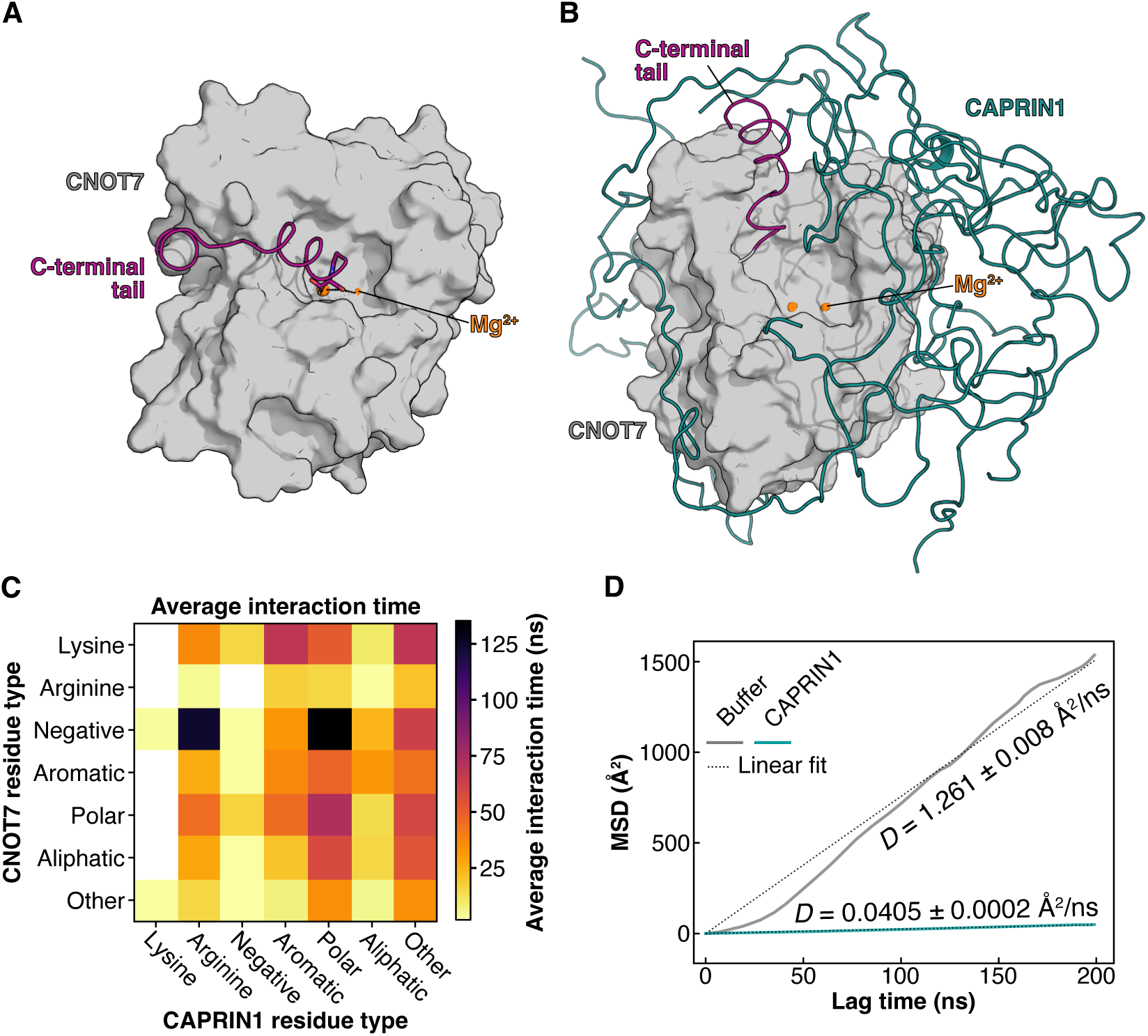
Solvation by CAPRIN1 modulates CNOT7 deadenylation activity. Final simulation frames of CNOT7 in (A) buffer and (B) 30 mM CAPRIN1 LCR (7 chains, teal) conditions, highlighting the C-terminal tail (pink) and Mg^2+^ ions (orange). (C) Average duration of intermolecular interactions (defined as ≤ 6 Å separating the centers of mass of two residues, one from each of CNOT7 and CAPRIN1), categorized by the ‘property’ of the residue at pH 7.4: lysine (K), arginine (R), negative (D, E), aromatic (F, H, W, Y), polar (N, Q, S, T), aliphatic (A, I, L, M, V), and other (C, G, P). White squares indicate ≤ 1 ns of interaction. (D) Mean squared displacement (MSD, Å^2^) of CNOT7 center of mass between simulation time *t* and time *t*+Δ*t* where Δ*t* is the lag time (ns) for CNOT7 in buffer (grey) or CAPRIN1 (teal) conditions with an open start pose. Diffusion coefficients (Å^2^/ns, listed on plot ± standard deviation) were determined from the slope of each MSD curve (eq S40). *R^2^* values for fits for buffer and CAPRIN1 are 0.976 and 0.985, respectively. See Figure S11B for full MSD plot (Δ*t* = 0-500 ns) and MSD data for closed start pose simulations.

The C-terminal tail of CNOT7 is predicted to be disordered (Figure S9A) with the potential to adopt both extended conformations as well as transient *α*-helical secondary structures. One (4GMJ) out of three deposited structures of CNOT7 in the RCSB PDB has partial electron density for the C-terminal tail docked in the active site. The others, 7AX1 and 7VOI, are missing tail density. In the 4GMJ structure^52^, the two active site Mg^2+^ ions coordinate E278 and E280 of the “EEE” motif (278-280) in the tail of CNOT7, with the tail in a relatively extended conformation (Figure 1A).

Based on our simulations starting from a helical IDPConformerGenerator-defined conformation (due to the helical propensity of the sequence, Figure S9A), we propose that the tail can also coordinate Mg^2+^ in a transient helical state through the backbone carbonyl and carboxyl functional groups of its last two residues (Q284 and S285). Note that IDPConformerGenerator uses a high-resolution non-redundant PDB database as the basis for torsion angle sampling, which constrains backbone geometry of the generated conformers to physically relevant values^54^. Interestingly, the AlphaFold2 (AF2) predicted structure for CNOT7 (AF2-Q9UIV1-F1-v6) shows the C-terminal tail in a low confidence conformation with helical structure overlapping the regions of the tail found to have higher helical propensity among the 10,000 IDPConformerGenerator models (Figure S9B). Additionally, the final pose from the buffer MD simulations is similar to the AF2 prediction (Figure S9B). The computations and simulations described above highlight the conformational heterogeneity possible for the disordered C-terminal tail of CNOT7 that can play a role in modulating the accessibility of RNA to the enzyme’s active site.

With the simulations in hand, we quantified pair-wise intermolecular interactions between residues in CNOT7 and CAPRIN1, focusing on the trajectories starting from an open pose, since the stability of the tail EEE-Mg^2+^ interactions and the absence of nearby negative counterions in the closed enzyme conformation likely impeded effective conformational sampling. Due to its highly positive net charge, we hypothesized that CAPRIN1 would bind the EEE motif in the CNOT7^WT^ tail to prevent it from occupying the active site. However, E278, E279, and E280 were among the residues with the fewest CAPRIN1 interactions, while Y227, S267, and Y268 of the tail had substantial contacts with several CAPRIN1 residues, primarily those that are aromatic and polar (Figure S10).

The MD reveals a multitude of solvating interactions between CAPRIN1 and the rest of CNOT7. Contacts with the longest average interaction times involved arginine-negative, polar-negative, aromatic-lysine, and polar-polar residues (Figure 5C). The impact of these solvating interactions can be observed in the dynamics of CNOT7. We tracked the position of CNOT7’s center of mass during the simulation (Figure S11A) and calculated its mean squared displacement (MSD) to estimate diffusion (Figures 5D and S11B). In both the open and closed starting pose simulations, CNOT7 diffusion was slower in the presence of CAPRIN1 (Figure 5D and S11B). Reduced diffusion rates in the condensate environment would impact enzymatic activity by impeding the rate of RNA-CNOT7 binding, in turn reducing catalytic efficiency.

We explored the potential for the solvating nature of CAPRIN1 condensates to be an evolutionarily conserved feature of the protein. While traditional multiple sequence alignment of IDRs usually does not identify sequence conservation^62^, the bulk sequence features of a disordered region (*e.g.*, amino acid content, charge, charge patterning, or aromatic patterning) are often conserved^63^ and can be associated with molecular function^64,65^. Using this approach (now referred to as FAIDR-ES: Feature Analysis of IDRs – Evolutionary Signatures), we found that the CAPRIN1 LCR shows evolutionary conservation of high tyrosine, phenylalanine and asparagine content, and the presence of RGG elements (Table S5), involved in phase separation and RNA binding^66,67^. The role of these amino acids in solvating CNOT7^WT^ suggests that they may play an important role in regulating cellular deadenylation.

## DISCUSSION

Our kinetic and microscopy data together with NMR and MD simulations provide a model for the regulation of CNOT7 activity in a CAPRIN1 condensate. The pre-equilibrium step in which CNOT7 exchanges between inactive and active conformations due to the disordered C-terminal tail either occluding or allowing binding of RNA to the enzyme’s active site, respectively, was confirmed to impede deadenylation (Figure 6A). We measured concentrations of RNA and enzyme in condensates and then quantified the extent to which the elevated concentrations lead to a reduction in the overall time required to degrade RNA (Figure 6B). The increased degradation from higher CNOT7 and RNA levels was countered by a decrease in catalytic efficiency, potentially originating from slower diffusion due to solvating interactions (*e.g*., electrostatics, hydrogen bonds, and cation-π interactions) between CAPRIN1 and CNOT7 (Figure 6C). Modified rates of catalysis could also contribute to the decrease in catalytic efficiency. In addition, the conformational equilibrium of CNOT7 was found not to be affected by the CAPRIN1 condensate environment (Figure 6D).

**Figure 6.**
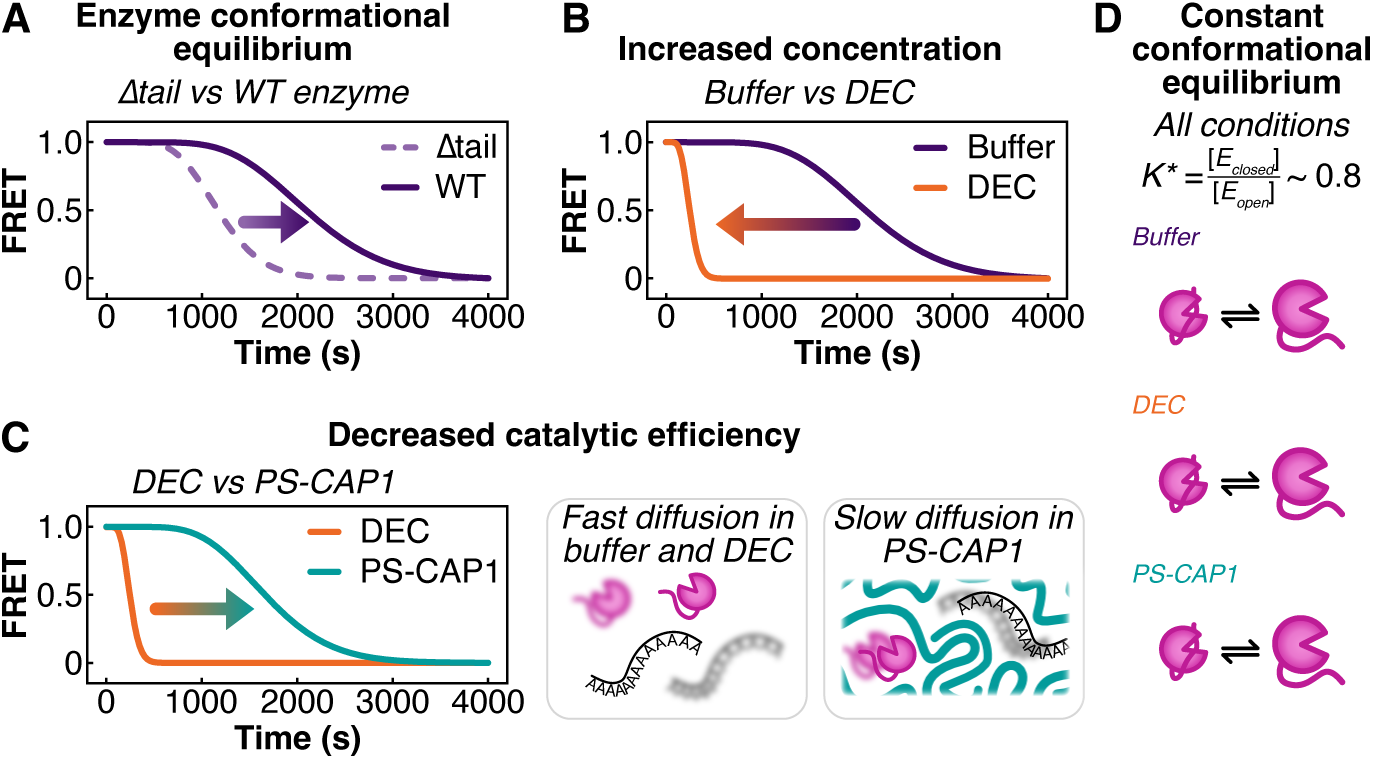
Visualization of the factors influencing deadenylation of RNA by CNOT7. (A) The CNOT7 tail significantly slows deadenylation. The FRET profile of deadenylation for a construct with an equilibrium between open/tail-free and closed/tail-bound conformations (WT, solid line) is shifted to the right (arrow) compared to a fully open construct (Δtail, dashed line). Effect is demonstrated with 3 µM enzyme, 100 nM RNA(A)_18_, *k_cat_*⁄*K_M_* = 3300 M^-1^s^-1^, and *K*^∗^ = 0 (Δtail) or *K*^∗^ = 0.8 (WT). (B) In the absence of hydrodynamic effects (e.g. slower diffusion) that offset concentration changes, increased concentrations of CNOT7 and RNA in the CAPRIN1 dense phase (24.3 µM enzyme, 442 nM RNA(A)_18_, orange) relative to buffer (3 µM bulk enzyme, 100 nM bulk RNA(A)_18_, purple) accelerate deadenylation. The FRET profile simulated with the enzyme and RNA DECs, representative of those in the condensates, is shifted left to earlier decay times as compared to buffer (arrow). The buffer and DEC curves were simulated with *k_cat_*⁄*K_M_* = 3300 M^-1^s^-1^ and *K*^∗^ = 0.8, values that approximate the catalytic efficiency and *K*^∗^ respectively in both cases (Table S1). (C) Catalytic efficiency of CNOT7 in PS-CAP1 condensates is diminished relative to DEC (24.3 µM enzyme, 442 nM RNA(A)_18_) in the absence of CAPRIN1. The simulated PS-CAP1 FRET profile shown here is shifted to the right (arrow) compared to the DEC curve resulting from the decrease in *k_cat_*⁄*K_M_* from 3300 M^-1^s^-1^ (DEC, orange) to 490 M^-1^s^-1^ (PS-CAP1, teal). Simulations for both conditions used *K*^∗^ = 0.8, which approximates the measured value in each case. Reduction in *k_cat_*⁄*K_M_* may be attributed, in part, to decreased diffusion in the dense phase, as was found in the molecular dynamics simulations. (D) The conformational equilibrium of the CNOT7 tail is unaffected by DEC or PS-CAP1 conditions (*K*^∗^ ∼ 0.8 in all cases) and therefore restricts deadenylation in these environments to the same extent as in buffer.

In addition to the effect of the CAPRIN1 condensate on deadenylation activity, which could occur directly through contacts with CNOT7 or by affecting the catalytic efficiency of the reaction, interactions between RNA and CAPRIN1 may also contribute significantly to changes in reaction rates. Hydrogen bonds, van der Waals interactions, and π-stacking of RNA ribose moieties or nucleobases with amino acid side chains contribute significantly to protein-RNA binding^68,69^. In particular, tyrosine, arginine, and phenylalanine (of which the 103-residue CAPRIN1 LCR has 7, 15, and 4, respectively) are known to form π-π interactions with RNA^68^. Indeed, single stranded RNA has been shown to diffuse and hybridize more slowly due to extensive interactions with CAPRIN1^60^. This would substantially slow the association of RNA with the active site and thereby reduce catalytic efficiency.

Biomolecular condensates can regulate enzymatic activity by shifting reactant concentrations and interaction networks, as shown here and elsewhere^27–32,35^. A notable recent example of this is the regulation of CCR4-NOT deadenylase activity by multivalent interactions with the Pumilio family RNA binding protein Puf3^70^. Similarly, decapping of mRNA by Dcp1/Dcp2 is autoinhibited by its own phase separation but reactivated upon incorporation of Edc3^35^. Multivalency of a condensate’s scaffold protein has further been shown to increase the enzyme E2 binding affinity for its substrate RanGAP^31^. It has also emerged that condensate environments can confer catalytic activity independent of enzymes^71,72^. Our study extends this growing body of work by providing detailed structural and kinetic insights into the mechanism underlying the regulation of CNOT7 activity by the CAPRIN1 condensate environment.

Studies of how ionic strength affects CAPRIN1 condensation have shown that the electrostatic potential, concentration, and hydrodynamic properties of its condensates are highly tunable^14,16,37^. Variations in condensate components could also impact the partitioning and diffusion of CNOT7 and RNA solvated within the condensate. Thus, shifting condensate composition could in principle provide a simple mechanism for modulating the magnitude of the different contributions to deadenylation and ultimately the rate of mRNA decay.

While we did not identify a shift in the conformational equilibrium of CNOT7 in the CAPRIN1 LCR condensate, this equilibrium significantly impacts CNOT7 activity and likely plays a role in regulating deadenylation by CCR4-NOT, consistent with previous suggestive work^56^. Disordered sequences may control enzymatic output through intermolecular interactions^73,74^, allosteric mechanisms^75,76^, or transient autoinhibition of active sites^77,78^. Furthermore, IDR elements are highly sensitive to their solvent environment and are therefore susceptible to regulation by condensates^79^. Biomolecular condensates are diverse and exhibit a wide range of solvent properties. As such, they are poised to tune conformational equilibria in general^14,22,26^, and, therefore, the activities of IDR-regulated enzymes.

Our quantitative methodology provides a physicochemical elucidation of changes in enzymatic activity in a biomolecular condensate environment. The complexity of biomolecular condensates allows them to synergize changes in biomolecule concentration, partitioning, and enzyme-substrate dynamics to enable sensitive control over biochemical reactions. The diversity of biomolecular condensates includes ones smaller than observable by light microscopy, likely including many RNA-protein complexes involved in RNA processing (*e.g*., splicing, translation, and degradation)^10^, extending the potential regulatory role of unique condensate environments with both protein and RNA contributions to the solvent. Future characterization of the multitude of the structural and chemical factors impacted by diverse condensate environments is needed to fully understand the functional regulation of enzymatic mechanisms.

## ASSOCIATED CONTENT

Supporting_information.pdf – contains materials and methods, Equations S1 to S40, Figures S1 to S12, Tables S1 to S4, Table S5 description

Table S5.csv – Evolutionary signatures of CAPRIN1 IDR2 (247-709) and LCR (607-709)

Zenodo repository – Microscopy data and analysis scripts, molecular dynamics trajectories and analysis scripts, and NMR data and analysis scripts can be found at doi:10.5281/zenodo.21080723

## Author Contributions

The manuscript was written through contributions of all authors. All authors have given approval to the final version of the manuscript. †These authors contributed equally.

## Funding Sources

Canadian Institutes of Health Research grant #PJT-190060 (JDFK)

Canada Research Chairs Program (JDFK)

National Institutes of Health grant R01GM127627 (THG, JDFK)

Natural Sciences and Engineering Research Council of Canada grant RGPIN-2024-05725 (JDFK), grant RGPIN-024-03872 (LEK), fellowship PGS-D 588933 2024 (ZHL)

SickKids Restracomp Fellowship (RWH)

## ACKNOWLEDGMENTS

The authors would like to thank Dr. Rashik Ahmed for fruitful discussions and valuable advice. They would also like to thank Dr. Jonathan Ditlev (The Hospital for Sick Children) and Gaddy Rakhaminov (The Hospital for Sick Children, University of Toronto) for help with experiments on the Elyra microscope, Dr. James Aramini for assistance at the University of Toronto Nuclear Magnetic Resonance Centre, Dr. Tae Hun Kim for acquiring and processing the CNOT7 NMR assignment measurements, both the SickKids Structural & Biophysical Core Facility and the SickKids Imaging Facility for the use of and guidance regarding their instruments.

## ABBREVIATIONS

AF2: Alpha Fold 2
CAPRIN1: Cytoplasmic Activation- and Proliferation-Associated Protein 1 (here, CAPRIN1 specifically refers to the LCR of full length CAPRIN1)
CI: confidence interval
DEC: droplet equivalent concentration
DVF: droplet volume fraction
FRET: Förster Resonance Energy Transfer
LCR: low complexity region
IDR: intrinsically disordered region
MD: molecular dynamics
NMR: nuclear magnetic resonance
PC: partition coefficient
PS-CAP1: phase-separated CAPRIN1
RNA(A)_18_: 5’ 6FAM-CCUUUCC(A)_18_
RGG: arginine-glycine-glycine
RNA(A)_38_: 5’ FITC-CCUUUCC(A)_38_
MSD: mean squared displacement;

## Notes

https://zenodo.org/records/21080723?token=eyJhbGciOiJIUzUxMiJ9.eyJpZCI6IjFiN2UyNDJlLWI3NzctNDgxMy04NjE3LWEyNDQwMzkwNDgzMCIsImRhdGEiOnt9LCJyYW5kb20iOiJlZTJmM2M3NDBiODkwYWU0ZWJhODc0MDg1OTZhZmVlZiJ9.rhmCa1aBrVrnD1ZRzYXfGhinwZrvAy9RhXD-6D112KaeXjujEuX0k-JajkGdowV88pHiXy-DOcmZfycFiDbydw

### Competing Interest Statement

The authors have declared no competing interest.

### Summary of Updates

The manuscript has not changed. The author list was updated to reflect the full list of authors.

