## Supporting information for "Competing effects modulate the rate of poly(A) RNA deadenylation in a biomolecular condensate"

The PDF file includes:

- Materials and Methods
- Figures S1 to S12
- Equations S1 to S40
- Tables S1 to S4
- Table S5 description

MATERIALS AND METHODS

Protein Purification

*$CNOT7^{WT}$ and $CNOT7^{\Delta tail}$ purification.* The gene for human CNOT7 ($CNOT7^{WT}$, 1-285) was previously inserted into a pET-SUMO vector[1]. The tailless CNOT7 ($CNOT7^{\Delta tail}$, 1-265, Figure S2) was generated by amplifying the region of interest from the pET-SUMO CNOT7 vector and using Gibson Assembly (New England Biolabs) to insert the gene into a linear pET-SUMO vector.

Each vector was transformed into *Escherichia coli* (*E. coli*) BL21-CodonPlus(DE3) RIPL cells and grown at 37 °C with kanamycin (50 μg/mL) in LB for assay and microscopy experiments or $^{15}N$-supplemented M9 minimal media[2] for NMR experiments. Protein expression was induced with 0.25 mM IPTG at an $OD_{600}$ ~ 0.8-1.0. Protein was expressed at 18 °C overnight and harvested via centrifugation at 4,000 × g for 20 min. The harvested cells were resuspended in lysis buffer (25 mM HEPES pH 7.8, 500 mM NaCl, 10 mM imidazole, 5 mM β-mercaptoethanol (BME), 2 mM $MgCl_2$, and 5% glycerol supplemented with DNase I, EDTA-free protease inhibitor cocktail (Sigma), and lysozyme).

Cells were lysed on ice using a Q Sonica Q500 (30% amplitude, 3 s on/5 s off, 7.5 min total time on). Debris was pelleted at 40,000 × g for 50 min and the supernatant was filtered through a 0.45 μm syringe filter and loaded on to ~20 mL of $Ni^{2+}$-NTA resin (Cytiva) that was pre-equilibrated with load buffer (25 mM HEPES pH 7.8, 500 mM NaCl, 10 mM imidazole, 5 mM BME, 2 mM $MgCl_2$, and 5% glycerol). The column was washed with 40 mL of load buffer and then 60 mL of wash buffer (load buffer with 30 mM total imidazole). The protein of interest was eluted with 60 mL elution buffer (load buffer with 300 mM total imidazole).

Eluted protein was treated with ULP1 protease (~25 μg/mL) to remove the $His_6$-SUMO tag and dialyzed against 25 mM HEPES pH 7.8, 300 mM NaCl, 5 mM β-mercaptoethanol (BME), 2 mM

$MgCl_2$, and 5% glycerol overnight at 4 °C. The tagless dialyzed protein was loaded onto the same Ni-NTA column re-equilibrated with load buffer and collected in the flow-through fraction. All steps up to this point were confirmed with SDS-PAGE.

The flow-through was concentrated to ~10 mL and run over a Superdex 75 26/600 column (Cytiva) equilibrated with dialysis buffer at 4 °C in 3.5 mL injections at 1 mL/min. Fractions were collected and purity was confirmed with SDS-PAGE and intact mass spectrometry.

*CAPRIN1 LCR purification.* The low-complexity region of CAPRIN1 (608-709) was prepared as described previously [1]. The vector for CAPRIN1 LCR or CAPRIN1 LCR with an additional C-terminal cysteine (C710) was transformed into *E. coli* BL21-CodonPlus(DE3) RIPL cells and grown at 37 °C with kanamycin (50 µg/mL) in 12-18 L of LB for assay and microscopy experiments. Protein expression was induced with 0.25 mM IPTG at an $OD_{600}$ ~ 0.8-1.0. Protein was expressed at 18 °C overnight and harvested via centrifugation at 4,000 × g for 20 min. The harvested cells were resuspended in lysis buffer (25 mM HEPES pH 7.8, 500 mM NaCl, 10 mM imidazole, and supplemented with DNase I, EDTA-free protease inhibitor cocktail (Sigma), and lysozyme; lysis buffer for CAPRIN1-Cys was supplemented with 5 mM BME).

Cells were lysed on ice using a Q Sonica Q500 (30% amplitude, 3 s on/5 s off, 2.5 min total time on). The slurry was then diluted 2-fold with 8 M guanidium chloride (GdmCl) and sonicated (30% amplitude, 3 s on/5 s off, 20 min total time on) with continuous stirring. Debris was pelleted at 40,000 × g for 50 min and the supernatant was filtered through a 0.45 µm syringe filter and loaded on to ~50 mL of Ni-NTA resin (Cytiva) that was pre-equilibrated with load buffer (25 mM HEPES pH 7.8, 4 M GdmCl, 500 mM NaCl, 10 mM imidazole, and 5 mM BME for the cysteine construct). The column was washed with 100 mL of load buffer and then 100 mL of wash buffer (load buffer with 30 mM total imidazole). Protein was eluted with 200 mL elution buffer (load buffer with 300 mM total imidazole).

Eluted protein was treated with ULP1 protease (~25 µg/mL) to remove the $His_6$-SUMO tag and dialyzed against 25 mM HEPES pH 7.8, 150 mM NaCl, 5 mM BME at 4 °C. Dialysis buffer was refreshed one time over 24 hours. The cleaved and dialyzed protein was diluted 2-fold with 8 M GdmCl and loaded onto the same Ni-NTA column re-equilibrated with load buffer and was subsequently collected in the flow-through fraction. All steps up to this point were confirmed with SDS-PAGE.

The flow-through was concentrated to ~10 mL and run over a Superdex 75 26/600 column (Cytiva) equilibrated with 25 mM Tris pH 8.0, 4 M GdmCl in 3.5 mL injections at 1 mL/min. Fractions were collected and purity was confirmed with SDS-PAGE and intact mass spectrometry.

<u>Fluorescent labeling</u>

Purified enzymes ($CNOT7^{WT}$ and $CNOT7^{\Delta tail}$) were labeled with Alexa Fluor 568 (AF568) or Alexa Fluor 647 (AF647), and CAPRIN1-Cys was labeled with AF647 (Invitrogen) via a maleimide linker. To label, the protein was incubated for 5 min with 5 mM DTT and then desalted over a 5 mL HiTrap Desalting column (Cytiva) into labeling buffer (CNOT7: 25 mM HEPES pH 7.2, 300 mM NaCl, 2 mM $MgCl_2$, and 5% glycerol, CAPRIN1: 25 mM HEPES pH 7.0, 4 M GdmCl). Fluorophore was added to ~200 µL of ~100 µM protein to a final concentration of 300 µM fluorophore. The protein-fluorophore mixture was put in a black, opaque microcentrifuge tube or a clear microcentrifuge wrapped with aluminum foil and left overnight at 4 °C on a rocker. The reaction was quenched via the addition of 5 mM DTT, and excess fluorophore and DTT were removed by desalting into assay buffer. Successful labeling was confirmed by intact mass spectrometry.

Experimental sample preparation

Prior to each experiment, purified proteins were buffer exchanged into assay buffers (given in following sections) via dialysis or by spin concentrator. Stock protein concentrations were measured at $A_{280}$ in triplicate with a NanoDrop One$^{C}$ (Thermo Scientific) before diluting to experimental concentration. These stocks were then diluted to the working concentrations given below.

**NMR spectroscopy**

Sample preparation

For measurements of $^{15}N$-$^{1}H$ HSQC spectra (Figure 1B), $^{15}N$-labeled CNOT7 was buffer exchanged by spin concentration into 50 mM sodium MES pH 6.5, 50 mM NaCl, 2 mM $MgCl_2$, 5 mM TCEP, and 10% $D_2O$/90% $H_2O$. $^{15}N$ CNOT7 samples for measurements of transverse relaxation rates, $R_2$ (Figure 1C), were prepared according to the following protocol. To remove trace background $Mg^{2+}$ in the buffer solutions for $R_2$ measurements, 1.1× NMR buffer base (55 mM sodium MES pH 6.5, 55 mM NaCl) was treated with Chelex 100 sodium form (Sigma, #95577) and stirred for 60 min, with pH correction to 6.5 after 30 min. A batch of 1× working NMR buffer (50 mM sodium MES pH 6.5, 50 mM NaCl, 10 mM TCEP, 10% $D_2O$) was then prepared with NMR buffer base, $D_2O$, and Chelex-treated TCEP pH adjusted to 6.5. Note that the $Mg^{2+}$ was first removed from these solutions with Chelex 100 and then added back to one of them using a known stock to better control the solution $Mg^{2+}$ concentration. CNOT7 samples were immediately buffer exchanged after purification by spin concentration into 1× working NMR buffer with 2 mM $MgCl_2$.

Data acquisition and analysis

All NMR spectra were acquired at 30 °C on an 18.8 Tesla (800 MHz $^1H$ frequency) Bruker Avance III HD spectrometer equipped with a cryogenically cooled x, y, z pulsed-field gradient triple-resonance probe. CNOT7 assignments were done using standard triple resonance approaches[3].

$^{15}N$ $R_1$ and $R_{1\rho}$ relaxation rates were measured for samples of 200 µM $^{15}N$-labeled CNOT7 with standard gradient enhanced sensitivity-based HSQC experiments[4]. A series of eight time points (5, 12.5, 20, 40, 60, 80, 100, and 120 ms) were collected for extraction of $R_{1\rho}$ values, while five time points (5, 150, 250, 400, and 600 ms) were measured for $R_1$ relaxation rates. The peak intensities as a function of relaxation time for the CNOT7 tail residues (266-285) were obtained using peakipy (https://j-brady.github.io/peakipy/). The relaxation profiles were fit with single exponential decays to obtain $R_{1\rho}$ and $R_1$ rates using an in-house Python script. $R_2$ values were then calculated using the relation

$$R_2 = (R_{1\rho} - R_1 \cdot \cos^2\theta)/\sin^2\theta \quad \text{(S1)}$$

where $\theta = \arctan(\nu_1/\Delta\Omega)$, $\nu_1$ is the spin-lock field strength (2 kHz), and $\Delta\Omega$ is the offset (Hz) of the spin in question from the $^{15}N$ carrier[5] .

**Deadenylation Assays**

Gel-based Deadenylation Assay (Figures 1 and S1)

FITC-labeled poly(A) RNA (5' FITC-CCUUUCC$(A)_{38}$, referred to as RNA$(A)_{38}$) was purchased as lyophilized samples from Sigma. 100 µM stocks were reconstituted in water and stored in -20 °C. Working stocks were diluted into reaction buffer (25 mM sodium phosphate pH 7.4, 100 mM NaCl, 2 mM $MgCl_2$, and 2 mM fresh DTT) upon use.

*In vitro* deadenylation assays with gel-based readout (Figures 1D and S1A) were adapted from Webster et al. (2017)[6]. In brief, CNOT7 or CNOT7$^{\Delta tail}$ (final concentration of 1.0 µM) was mixed with varying concentrations of RNA$(A)_{38}$ in a reaction buffer. Reaction mixtures were incubated at 30 °C. Samples of the reactions (5 µL) were removed and quenched at specific time points by mixing in a 2× denaturing loading dye (95% formamide, 10 mM EDTA, 0.01% w/v bromophenol blue). Samples were boiled for 5 min before loading onto a 15% TBE-Urea gel (Thermo Scientific). Gels were run in 1× TBE buffer for 110 min at a constant voltage of 100 V. After running, gels were directly scanned with a GE Typhoon FLA9500 Phosphorimager (GE Healthcare) to detect FITC signal and RNA$(A)_{38}$ migration (Figure S1A).

Gel images were analyzed using Fiji and Python to calculate deadenylation rates based on displacement of cleaved RNA$(A)_{38}$ as a function of time (Figure S1B). A manually centered, straight, vertical line was drawn from edge to edge of the image over each lane. The lane profiles were fit with a multi-gaussian function to find the peak with the maximum volume, which was assumed to represent the average distance travelled by the poly(A) RNA product distribution at each time point. The difference between the major peak position for the time t = 0 s sample (no poly(A) decay) and those of each later timepoint were plotted against time and fit with a straight line to calculate an apparent rate of degradation (Figure S1B). Data from lanes for which the reaction was complete were not included in fits. The slopes of these lines, $y$, were plotted against substrate concentration, $[RNA]$ (Figure S1C). Note that it is not straightforward to use the

Michaelis-Menten model to fit these data as the rates are calculated in terms of distances travelled in the lanes of the gels, not product concentrations[6].

FRET-based Deadenylation Assay (Figure 2)

6FAM-labeled poly(A) RNA (5' 6FAM-CCUUUCC$(A)_{18}$, referred to as RNA$(A)_{18}$), and TAMRA-labeled poly(T) DNA (5' $(T)_{38}$GGAAAGG-TAMRA, referred to as DNA$(T)_{38}$), were purchased as lyophilized samples from Sigma. Tubes were stored at -20 °C until use, when one stock was reconstituted to 100 μM in RNase-free water (from an EMD Millipore Advantage A10 water system equipped with a Biopak Polisher). CNOT7 enzyme was dialyzed over night at 4 °C into assay buffer (50 mM sodium phosphate pH 7.4, 50 mM NaCl, 2 mM $MgCl_2$, and 5 mM fresh DTT). CAPRIN1 was dialyzed for at least 48 hours at room temperature in 25 mM sodium phosphate pH 7.4. During this time, CAPRIN1 dialysis buffer was refreshed 3 times at evenly spaced intervals.

The deadenylation assay, originally adapted from Maryati, *et al.* (2014)[7], was run as described previously[8] in a Synergy Neo2 plate reader equipped with an automated dual-injector system[8]. Aliquots of enzyme (1-7 μM for buffer and PS-CAP1 experiments, see Table S3 for DEC experimental enzyme concentrations) were distributed down a column in a 384-well plate (Figure 2). All experiments included a 0 μM enzyme condition for background correction. Using the injectors to ensure rapid and consistent dispensation, DNA$(T)_{38}$ (at 5× the concentration of RNA) in quenching buffer (50 mM sodium phosphate pH 7.4, 50 mM NaCl, 5 mM fresh DTT, 5 mM EDTA, 0.5% SDS, and 0.1 mg/ml Proteinase K – concentrations reflect final concentrations in the well) was added to the first row for the $t = 0$ points (Figure 2). After 1 minute of shaking, RNA$(A)_{18}$ was added to each well (5 μL at 500 nM for low concentration experiments or 2.18 μM for DEC experiments) to achieve a final concentration of 100 nM for the 1-7 μM enzyme concentrations or 436 nM for DEC experiments. Rows were sequentially quenched over the course of ~5-60 min with DNA(T)$_{38}$ in quenching buffer. The plate was continuously shaking between quenches.

Degradation of the substrate by CNOT7 generates species of varying poly(A) length (1-18 bases), each of which has a unique binding affinity with the DNA(T)$_{38}$. The 5' CCUUUCC and 3' GGAAAGG motifs in RNA(A)$_{18}$ and DNA(T)$_{38}$ respectively act as annealing tags that orient the oligomers. The complementary tags eliminate different registers of the poly(A) tail-poly(T) trap strand binding and ensure that the fluorophores are proximal upon annealing, allowing for resonance energy transfer between them. Shorter poly(A) RNA species have a lower binding affinity with DNA(T)$_{38}$ (see *Deadenylation kinetic model: RNA-DNA annealing and fitting the FRET profiles*), and as these shorter species are generated, the FRET between the 6FAM and TAMRA fluorophores decreases due to less annealing. The CCUUUCC sequence in the RNA strand also serves as a form of 3'-UTR, which would be present in cellular mRNA.

Fluorescence was measured at excitation/emission wavelengths 485/528 ($I_{DD}$), 485/590 ($I_{DA}$), 550/590 ($I_{AA}$) with a spectral width of 10 nm and gain at 80. $I_{DD}$ was normalized by subtracting the $I_{DD}$ from the 0 μM enzyme condition, and $I_{DA}$ was normalized at each timepoint by dividing $I_{DA}$ by the ratio of the $I_{AA}$ at that timepoint to the $I_{AA}$ at time t = 0 s. FRET was then calculated as

$$FRET = \frac{I_{DA}}{I_{DD}+I_{DA}} \tag{S2}$$

FRET deadenylation assays were run multiple times at 25 °C in buffer ("buffer conditions", five replicates), with CAPRIN1 ("PS-CAP1 conditions", five replicates), and at the droplet equivalent concentration of enzyme and RNA ("DEC conditions", four replicates). See exemplary data in Figure 4A and remaining data in Figure S4. FRET data were fit as described below.

The time required to reach FRET = 0, $t_{F=0}$, was approximated by eye for all CNOT7$^{WT}$ experiments. These times were reported as averages ± 1 standard deviation.

**Deadenylation kinetic model**

The kinetic model used to fit our FRET data can be written as

$$E_{closed} \underset{k_{-1}}{\overset{k_1}{\rightleftharpoons}} E_{open} + TA_i \underset{k_{-2}}{\overset{k_2}{\rightleftharpoons}} ETA_i \xrightarrow{k_{cat}} E_{open} + TA_{i-1} + A_1 \quad \text{(S3)}$$

where $k_1$ and $k_{-1}$ are the rates of interconversion between the inactive (closed) enzyme ($E_{closed}$) and the active (open) enzyme ($E_{open}$), $k_2$ is the rate of association of free RNA ($TA_i$) with active enzyme, $k_{-2}$ is the corresponding dissociation rate of enzyme-RNA complexes ($ETA_i$), and $k_{cat}$ is the catalytic rate for the reaction whereby a single adenosine ($A_1$) is released from the substrate, producing a shorter free RNA product ($TA_{i-1}$), and free active enzyme. For simplicity, we assume that $k_2$, $k_{-2}$, and $k_{cat}$ are the same for poly(A) RNA of any length. Note that the poly(A) RNA substrate annealing tag is denoted as $T$.

Generation of RNA(A) subspecies

Based on eq S3 (Scheme 1 in main text), the concentration of each RNA species $TA_i$ at time, *t*, is given by

$$[TA_i]_{total} = [TA_i] + [ETA_i] \quad \text{(S4)}$$

and their time dependencies can be described according to

$$\frac{d}{dt}[TA_i]_{total} = \frac{d}{dt}[TA_i] + \frac{d}{dt}[ETA_i] \quad \text{(S5)}$$

If we assume, for the moment, that $[E_{closed}] = 0$ (*i.e.*, the enzyme is fully open as is the case for CNOT7$^{\Delta tail}$), and, further, that species $TA_1$ is not hydrolysable, we can describe the change in RNA species as

$$\frac{d}{dt}[TA_n] = -k_2[E_{open}][TA_n] + k_{-2}[ETA_n] \quad \text{(S6)}$$

$$\frac{d}{dt}[TA_i] = -k_2[E_{open}][TA_i] + k_{-2}[ETA_i] + k_{cat}[ETA_{i+1}], 1 \le i < n \quad \text{(S7)}$$

$$\frac{d}{dt}[ETA_i] = k_2[E_{open}][TA_i] - k_{-2}[ETA_i] - k_{cat}[ETA_i], 2 \le i \le n \quad \text{(S8)}$$

$$\frac{d}{dt}[ETA_1] = k_2[E_{open}][TA_1] - k_{-2}[ETA_1] \quad \text{(S9)}$$

which results in the following when combined with eq S5:

$$\frac{d}{dt}[TA_n]_{total} = -k_{cat}[ETA_n] \quad \text{(S10)}$$

$$\frac{d}{dt}[TA_i]_{total} = k_{cat}([ETA_{i+1}] - [ETA_i]), 2 \le i < n \quad \text{(S11)}$$

$$\frac{d}{dt}[TA_1]_{total} = k_{cat}[ETA_2] \quad \text{(S12)}$$

where $n$ = 18 in our case (*i.e.*, the RNA substrate contains an 18-adenosine tail). To reduce the parameter space (from five kinetic parameters to one) and obtain well defined constants for comparing the CNOT7 constructs we used, we implemented the steady state assumption (justified below in *'Evaluating the steady state assumption and assessing the ratio of $E_{closed}$ and $E_{open}$'*, Figure S3),

$$\frac{d}{dt}[ETA_i] \approx 0 \quad \text{(S13)}$$

along with the following definition for the Michaelis constant,

$$K_M \equiv \frac{k_{-2}+k_{cat}}{k_2} \quad \text{(S14)}$$

and the assumption that $[ETA_i]$ is negligible compared to $[E]_{total}$ because our assays were performed under conditions where $[E]_{total} \gg [TA_i]$ so that $[E_{open}] \sim [E]_{total}$. This results in a simplification of eqs S10-S12 to

$$\frac{d}{dt}[TA_n]_{total} = -k_{obs}[E]_{total}[TA_n] \quad \text{(S15)}$$

$$\frac{d}{dt}[TA_i]_{total} = k_{obs}[E]_{total}([TA_{i+1}] - [TA_i]),\ 2 \le i < n \quad \text{(S16)}$$

$$\frac{d}{dt}[TA_1]_{total} = k_{obs}[E]_{total}[TA_2] \quad \text{(S17)}$$

where $k_{obs}$ is the apparent rate for a fully open enzyme (*i.e.*, $k_1 >> k_{-1}$, represented experimentally by CNOT7$^{\Delta tail}$), defined as

$$k_{obs} = \frac{k_{cat}}{K_M} = \frac{k_{cat}}{\frac{k_{-2}+k_{cat}}{k_2}} \quad \text{(S18)}$$

For CNOT7$^{WT}$, where $[E_{closed}] \neq 0$, we define $[E]_{total}$ as

$$[E]_{total} = [E_{closed}] + [E_{open}] + \sum_{i=1}^{n}[ETA_i] \quad \text{(S19)}$$

Assuming a pre-equilibrium between closed ($E_{closed}$) and open ($E_{open}$) tail conformations (see *'Evaluating the steady state assumption and assessing the ratio of* $E_{closed}$ *and* $E_{open}$*'* below for justification, Figure S3) such that

$$K^* = \frac{[E_{closed}]}{[E_{open}]} = \frac{k_{-1}}{k_1} \quad \text{(S20)}$$

and, once again, assuming $[ETA_i]$ is negligible relative to $[E_{open}]$, we arrive at the following definition for $[E]_{total}$

$$[E]_{total} = K^*[E_{open}] + [E_{open}] \quad \text{(S21)}$$

which allows us to modify eqs S15-S17 and calculate the change in RNA concentrations in the presence of CNOT7$^{WT}$ according to

$$\frac{d}{dt}[TA_n]_{total} = -k_{obs}^*[E]_{total}[TA_n] \quad \text{(S22)}$$

$$\frac{d}{dt}[TA_i]_{total} = k_{obs}^*[E]_{total}([TA_{i+1}] - [TA_i]),\ 2 \le i < n \quad \text{(S23)}$$

$$\frac{d}{dt}[TA_1]_{total} = k_{obs}^*[E]_{total}[TA_2] \quad \text{(S24)}$$

where the effective rate constant optimized in the data fitting routine, $k_{obs}^*$ (Figures 4B and S5, Table S1), is defined as

$$k_{obs}^* = \frac{k_{cat}}{K_M(1+K^*)} = \frac{k_{cat}}{\frac{k_{-2}+k_{cat}}{k_2}(1+\frac{k_{-1}}{k_1})} \tag{S25}$$

The above derivation points to a simple way of isolating the conformational equilibrium constant, $K^*$ (Figure 4C), from fitted $k_{obs}$ (FRET data recorded using CNOT7$^{\Delta tail}$) and $k_{obs}^*$ (FRET data recorded using CNOT7$^{WT}$), via the relation

$$K^* = \frac{k_{obs}}{k_{obs}^*} - 1 = \frac{[E_{closed}]}{[E_{open}]} \tag{S26}$$

which assumes that the presence of the tail in the open form or the absence of the tail does not impact substrate binding or catalysis. The fraction tail closed and open in the overall CNOT7$^{WT}$ population (Figure 4D) are then calculated according to

$$Frac_{closed} = \frac{K^*}{1+K^*} \ , \ \ Frac_{open} = \frac{1}{1+K^*} \tag{S27}$$

To fit the FRET data to obtain $K^*$ values, it is first necessary to recast $[TA_i]$ in terms of measured FRET values, which can be accomplished as described briefly immediately below, and in more detail previously[8].

For a given experiment, assays with CNOT7$^{WT}$ and CNOT7$^{\Delta tail}$ were run in parallel and the $K^*$ value for CNOT7$^{WT}$ was calculated from the resulting $k_{obs}^*$ and $k_{obs}$. The $k_{obs}^*$ values as well as the $K^*$ values reported in Figure 4B and Table S1 are the average of 4 or 5 experiments. See Figure 4A and Figure S4 for all experimental data and see Figure S5 for individual experiment fit results. Note that in the main text we refer to both $k_{obs}^*$ and $k_{obs}$ as $k_{obs}^*$, for simplicity, but it is understood that the two are equivalent only in the limit that $K^*$ = 0, that is, when the enzyme is in the fully open or state (or for CNOT7$^{\Delta tail}$).

<u>RNA-DNA annealing and fitting the FRET profiles</u>

Experimentally, the deadenylation reaction is quenched at regular time points by adding 1% SDS, 0.1 mg/mL Proteinase K, and a 3' TAMRA fluorophore-conjugated DNA strand (5' $(T)_{38}$GGAAAGG-TAMRA, referred to as DNA$(T)_{38}$) that is complementary to the full-length RNA substrate and the deadenylation reaction intermediates. The time-dependent FRET signal is then recorded. Computationally, this is modeled by explicitly calculating values for $[TA_i]_{total}$ via numerical integration of eqs S22-S24 at each timepoint $t$. The distribution of RNA-DNA hybrid duplexes, $TA_iD$, is then obtained from a system of equations constrained by mass conservation (*i.e.,* the total added DNA and the set of $[TA_i]_{total}$ are given by sums of free and annealed concentrations) and the relations

$$[TA_iD] = [TA_i][D]K_{TA_iD} = [TA_i][D]e^{\frac{-\Delta G_{TA_iD}}{RT}} \quad \text{(S28)}$$

where $TA_iD$ is RNA species $TA_i$ annealed to DNA, $D$ is free DNA, $R$ is the ideal gas constant, and $T$ is the experimental temperature in Kelvin. In eq S28, the free energy of annealing between each RNA species $TA_i$ and DNA$(T)_{38}$, $\Delta G_{TA_iD}$, was calculated according to:

$$\Delta G_{TA_iD} = \alpha \cdot i + \Delta G_0 \quad \text{(S29)}$$

where $i$ is the number of adenosine bases, $\alpha$ is the annealing scaling parameter, and $\Delta G_0$ is the free energy of binding between the RNA and DNA annealing tags. In the analysis of the deadenylation data recorded under buffer or DEC conditions, the annealing parameters were calculated with nearest-neighbor parameters for RNA-DNA hybridization[9], resulting in an $\alpha$ value of -4.0684 kJ/mol and $\Delta G_0$ value of -3.2498 kJ/mol. Analysis of the profiles generated under the PS-CAP1 condition involved an additional grid-search step where the value of $\alpha$ was optimized to minimize the fit RMSD (Figure S12). The resulting $\alpha$ value used in PS-CAP1 fits was -2.7351 kJ/mol.

The population of each RNA species $TA_i$ bound to DNA, $P_{TA_iD}$, was calculated according to

$$P_{TA_iD} = \frac{[TA_iD]}{[TA_n]_{t=0}} \tag{S30}$$

where $TA_n$ is the total concentration of RNA substrate at time $t$ = 0. Once $P_{TA_iD}$ values are available the expected FRET at a given timepoint, $F_{calc,t}$, is calculated as

$$F_{calc,t} = \Delta F \sum_{i=1}^{n} P_{TA_iD} + F_o \tag{S31}$$

where $\Delta F$ is the range of FRET values in the kinetic profile (*i.e.*, from $t$ = 0 to $t \to \infty$), $F_o$ is the baseline FRET value ($t \to \infty$), $n$ is the number of adenosine residues in the full-length RNA substrate (18 in this case), and $i$ is the length of each RNA degradation product. Fits were carried out by adjusting $k^*_{obs}$, $\Delta F$, and $F_o$ to minimize the residual sum of squares between experimental FRET points and $F_{calc,t}$.

For information regarding the fit of phase-separated CAPRIN1 deadenylation data, refer to the *'Fitting PS-CAP1 deadenylation assay data with dense-phase concentrations'* section below.

Determining the accuracy of the rate constant $k^*_{obs}$ and the equilibrium constant $K^*$ (Figure 3)

To assess whether it is possible to reliably extract meaningful parameters from our experimental FRET data using our kinetic model, we performed a systematic validation with simulated datasets, illustrated in Figure 3. Although the legend to this figure summarizes what was done, we elaborate on some of the details in this section. Our simulated datasets were leveraged in fitting routines to explore whether the fitted rate constants, and the $K^*$ and $k^*_{obs}$ calculated from them, could recapitulate the "ground truth" values used to generate the synthetic FRET data. We simulated time-course data for RNA deadenylation starting from several initial conditions of the CNOT7 tail equilibrium that represent different proportions of enzyme in the "open" (catalytically competent) *vs* "closed" (incompetent) conformational states (Figure 3A). Values for the 5 rate constants in the model ($k_1$, $k_{-1}$, $k_2$, $k_{-2}$, and $k_{cat}$) were constrained so that the $k^*_{obs}$ values calculated from them were distributed about the experimental $k_{obs}^{*}$ value measured for CNOT7 in buffer at 25 °C ($1.82\times10^{3}$ $M^{-1}$ $s^{-1}$). The value for $k_1$ was set to 1 $s^{-1}$ while $k_{-1}$ assumed a range of values, $5.26\times10^{-2}$, $2.5\times10^{-1}$, 1, 4, and 19 $s^{-1}$, to generate simulated data for 5, 20, 50, 80, and 95% closed CNOT7 at equilibrium, respectively. Values for $k_2$, $k_{-2}$, and $k_{cat}$ were set to $1\times10^{5}$ $M^{-1}$ $s^{-1}$, 3 $s^{-1}$, and $1.15\times10^{-1}$ $s^{-1}$ respectively, for all simulated datasets to yield FRET progress curves that were similar to the ones measured. This generated a set of $k_{obs}^{*}$ values ($3.51\times10^{3}$, $2.95\times10^{3}$, $1.85\times10^{3}$, $7.38\times10^{2}$, $1.85\times10^{2}$ $M^{-1}s^{-1}$) for fraction closed CNOT7 of (5, 20, 50, 80, 95%). The synthetic FRET curves for each fraction closed CNOT7 were calculated using 1, 2, 3, 5, 7, and 10 μM total enzyme with 0.1 μM $RNA(A)_{18}$ (Figure 3A), in keeping with our experimental conditions. Gaussian noise was added to these datasets using a normal distribution centered at 0 and a standard deviation equal to a root mean squared deviation (RMSD) of $5.555\times10^{-2}$ that was obtained in fits of our data.

In our analysis we tested whether the fitting algorithm converged to the ground truth parameters that were used to generate the synthetic datasets (see above), regardless of the values of the starting fit parameters. To test for extraction of the ground truth parameters and consistent convergence of the fits, we selected the synthetic 50% closed CNOT7 dataset as a proxy for the experimental data in buffer. We used Latin hypercube sampling (LHS) from the *SciPy* Python package[10] to draw starting fit parameter sets over wide ranges. LHS randomly samples parameters from multidimensional parameter spaces with better coverage and spacing than simple random sampling. This reduces clustering or gaps in parameter samples that might otherwise occur and allows for a dramatic reduction of the computation time required to explore wide input parameter ranges in fits of models with multiple parameters. Parameter ranges for LHS were chosen to span several orders of magnitude – $k_1$: [$1\times10^{-2}$, $1\times10^{2}$] $s^{-1}$, $k_{-1}$: [$1\times10^{-2}$, $1\times10^{2}$] $s^{-1}$, $k_2$: [$1\times10^{4}$, $1\times10^{7}$] $M^{-1}$ $s^{-1}$, $k_{-2}$: [$1\times10^{-1}$, $1\times10^{2}$] $s^{-1}$, and $k_{cat}$: [$1\times10^{-3}$, $1\times10^{1}$] $s^{-1}$. In total, 200 initial parameter sets were used as inputs to fit the 50% closed dataset, from which 200 output parameter sets were obtained. Of the 200 inputs, 179 led to high quality fits (RMSD < $7\times10^{-2}$). These input parameter sets and their corresponding outputs were selected for further analyses (Figure 3B). The remaining output parameter sets were discarded as they were derived from inputs that were too far from optimal, leading to poor minimization by the fitting routine (no convergence).

For the selected starting fit parameter sets, we explored how strongly they were correlated with the output values, and most importantly whether the outputs reliably reproduced the ground truth values used to simulate the synthetic 50% closed dataset we fit. We found that the individual rate constants extracted from the fits were in general not consistent with the ground truth values. Instead, they were correlated with the initial values (Figure 3B), indicative of large rate constant covariances during minimization. Some of the fitted $k_{cat}$ values, however, did match the ground truth when initiated over the range of ~$10^{-3}$ to $10^{0}$ $s^{-1}$, highlighting that the fit routine is at least partly sensitive to $k_{cat}$. Interestingly, when $k_{cat}$ was initiated above roughly $10^{0}$ $s^{-1}$, it did not adjust to the ground truth but was instead correlated with its input value like the other parameters. As anticipated from these results, the $K^*$ generated from the fitted $k_1$ and $k_{-1}$ ($K^* = k_{-1}/k_1$) also did not recapitulate the ground truth $K^*$ and was correlated with that calculated from the inputs (Figure 3B). We subsequently calculated $k^*_{obs}$ from the relation $k^*_{obs} = k_{cat}/K_M(1 + K^*)$ (see eq S25 for derivation; eq 1 in main text). Even with starting fit parameters that resulted in starting $k^*_{obs}$ rates spread widely over the $10^{-1}$ to $10^{6}$ $M^{-1}$ $s^{-1}$ range, the fitted $k^*_{obs}$ values were tightly clustered around the ground truth value (Figure 3C), and, in turn, the $K^*$ calculated using the fitted $k^*_{obs}$ and the ground truth $k_{obs}$ accurately reproduced the ground truth $K^*$ value. Thus, although individual rate constants cannot be determined accurately, robust values of $k^*_{obs}$ are generated. To further test this, we subsequently extended our analysis to the additional $K^*$ values, corresponding to 5%, 20%, 80%, and 95% closed. In all cases, fitted $k^*_{obs}$ and $K^*$ were tightly distributed about the values calculated from the ground truth parameters used to simulate the data (Figure 3D and 3E), providing confidence that these parameters can be robustly extracted from fits to our FRET data.

Evaluating the steady state assumption and assessing the ratio of $E_{closed}$ and $E_{open}$

Our kinetic model is based on the steady state assumption whereby $\frac{d}{dt}[ETA_i] \approx 0$, and holds for the case where $[ETA_i] << [E]_{Total}$, which is relevant for our experimental conditions. The $k^*_{obs}$ values extracted from our fits are a function of $K^*$, and hence the concentrations of $E_{closed}$ and $E_{open}$ at equilibrium. In order to establish how the concentrations of enzyme-bound RNA, $E_{closed}$, and $E_{open}$ vary during the course of the deadenylation reaction, we have carried out a set of simulations using the same kinetic parameters as in the section immediately prior (a parameter table is given in Figure S3A), except with $k_1$ and $k_{-1}$ both set to either $1\times10^{-2}$ (slow equilibration of $E_{closed}$ and $E_{open}$) or $1\times10^{2}$ $s^{-1}$ (rapid equilibration), corresponding in both cases to $K^*$ = 1 (50% closed), Figure S3B and S3C, respectively.

In both cases we found that the sum of the reaction rates for the RNA-bound enzyme species (black lines) remained constant or very near constant for the bulk of the reaction (*i.e.*, in steady state) due to the sequential nature of the deadenylation mechanism and that the binding, dissociation, and catalytic rate constants are independent of RNA size (Figure S3B and S3C; colored curves correspond to bound RNAs of different lengths, decreasing in size from red to blue). Only once much smaller poly(A) species (roughly < $A_{10}$) are produced and subsequently degraded is there observable decay of RNA-bound enzyme, evidenced by the small fluctuations in the reaction rates. This is only visible with the 10 μM enzyme concentration where the trajectories go to completion for the reaction parameters considered here (Figure S3B and S3C, right). However, because these concentrations are very small to begin with they essentially do not change throughout the course of the reaction trajectory, validating the steady state assumption.

With either slow or fast equilibration of $E_{closed}$ and $E_{open}$ at either 1 or 10 µM CNOT7 and 0.1 µM RNA(A)$_{18}$ (our assay concentrations), we observed effectively no deviation from the equilibrium unbound enzyme concentration values ($d[E_{open}]/dt = d[E_{closed}]/dt = 0$ orange- $[E_{closed}]$ and green- $[E_{open}]$ lines, subpanels in Figure S3B and S3C where the derivatives have been normalized to the total enzyme concentration), as there is not enough RNA to influence the enzyme pool dramatically.

Evaluating changes in catalytic efficiency

$k^*_{obs}$, the apparent rate of deadenylation ($M^{-1}s^{-1}$; eq S25), is derived from two components, $k_{obs} = k_{cat}/K_M$ (eq S18), and $K^*$ (eq S20), the former of which is sensitive to the properties of the solvent in which the reaction occurs, while the latter reports on the conformational equilibrium of the tail. An important goal is to compare $k^*_{obs}$ values obtained under PS-CAP1 or DEC conditions with the corresponding rates measured in buffer to evaluate how changes in $K^*$ impact the observed rates. It follows directly from eq S25 that

$$\frac{k_{obs,i}}{k_{obs,j}} = \frac{(1+K^*_j)}{(1+K^*_i)} \cdot \frac{k^*_{obs,i}}{k^*_{obs,j}} \qquad \text{(S32)}$$

where $K^*_i$ and $k^*_{obs,i}$ or $K^*_j$ and $k^*_{obs,j}$ are measured under two different conditions, $i$ and $j$ (Figure 4E, Table S4). In the case of CNOT7$^{\Delta tail}$, $k_{obs} = k^*_{obs}$ because $K^* = 0$. Table S1 lists $k^*_{obs,i}$ and $K^*_i$ values for all the conditions examined in this work.

### Confocal Microscopy

Instrumentation and sample preparation for colocalization experiments

Microscopy was performed with the same 6FAM-RNA$(A)_{18}$ and buffer and sample volumes as used in the FRET-based deadenylation assays. For colocalization experiments involving CNOT7, RNA, and CAPRIN1, 6FAM-RNA$(A)_{18}$ (100 nM), AF568-labeled enzyme (total 1 or 10 µM enzyme, 50 nM labeled), and/or CAPRIN1 (100 µM total CAPRIN1, 50 nM labeled) were combined depending on sample components or replaced with the equivalent volume of buffer. Components were mixed such that unlabeled CAPRIN1 was added second to last and 6FAM-RNA$(A)_{18}$ (or buffer for controls) was added immediately prior to imaging. The 25 µL sample was transferred to a 96-well black cell imaging plate with a glass bottom (Eppendorf #0030 741.030).

Fluorescence images to observe colocalization of CNOT7 or CNOT7$^{\Delta tail}$ with CAPRIN1 and RNA$(A)_{18}$ were acquired on a Leica DMi8 spinning disk confocal microscope equipped with a 63×/1.3 NA oil immersion objective, a Hamamatsu C9100-13 EM-CCD camera, and Spectral Borealis lasers. Images were acquired with 488 nm/525 nm (50 nm SW), 561 nm/600 nm (60 nm SW), and 637 nm/700 nm (75 nm SW) excitation/emission channels at ambient temperature (Figure S6).

Instrumentation and sample preparation for experiments to determine droplet equivalent concentration (DEC)

Samples of CNOT7 or CNOT7$^{\Delta tail}$ (50 nM AF647-labeled enzyme, total 0-5 µM), ± CAPRIN1 (100 µM unlabeled) and ± 6FAM-RNA$(A)_{18}$ (100 nM) were combined in a similar manner as the colocalization experiments. Samples were imaged at ambient temperature in a PF127-passivated 96-well glass bottom plate[11], which helped reduce background noise and increase droplet resolution. Six samples were prepared at a time. Immediately prior to imaging of samples, the proteins were dispensed into the imaging plate. RNA$(A)_{18}$ was then added, and the plate was centrifuged at 500 × g for 30 s at 25 °C to encourage drops to settle to the base of the wells.

Calibration curves were generated by imaging titrated samples of 6FAM-RNA$(A)_{18}$ (20-800 nM), AF647-CNOT7 (0.1-2.5 μM), and AF647-CNOT7$^{\Delta tail}$ (0.1-2.2 μM) in assay buffer.

Image acquisition

Images for the DEC experiments were acquired with a Zeiss Elyra PS.1 microscope (Zeiss) equipped with a 100×/1.46 NA oil immersion objective, Andor iXon 897 EMCCD camera, and 488 and 647 nm HR-Diode lasers. Zen software (Zeiss) was used for image acquisition.

For each sample, the focal plane was centered just above the bottom of the well. To acquire dilute-phase and bulk (non-phase-separated condition) images, the focal plane was moved at least 100 μm up from the well base where the sample was almost or completely homogeneous (*i.e.*, no droplets in frame). Droplets were imaged by returning the focal plane to the bottom of the well, selecting a pixel in a background (non-droplet) region of the live image, and then raising the focal plane such that the droplets were still in the focal plane but the background intensity was close to that of the dilute-phase images (*i.e.*, the point at which the imagine plane was approximately at the equator of the droplets of interest) (Figure S7A).

Image analysis

Microscopy images were exported from Volocity or Zen as stacked, 512 × 512-pixel TIFF files and analyzed in an automated fashion with Fiji[12] and Python scripts (provided in the supplemental archive). In brief, images were cropped to a 384 × 384-pixel square in the center of the image. For each channel, a histogram of 256 bins was generated. The thresholds for the foreground, $T_f$, and background, $T_b$, were empirically determined according to

$$T_f = I_{max} + 3(I_{max} - I_{min}) \quad \text{(S33)}$$

$$T_b = I_{max} + 1.5(I_{max} - I_{min}) \quad \text{(S34)}$$

where $I_{max}$ is the intensity value for the bin with the highest pixel count and $I_{min}$ is the lowest intensity value of the image. Pixels of intensity equal to or higher than $T_f$ were counted as "in-drop" pixels. Pixels of intensity equal to or lower than $T_b$ were counted as "out-of-drop" pixels. The thresholds were used to generate foreground and background masks, which were used to delineate regions inside and outside drops.

For droplet equivalent concentration experiments, particles in the foreground mask were clarified with the "Adjustable Watershed" Fiji plugin[13] and identified with the "Analyze particles" Fiji tool. For each particle larger than 6 pixels with an aspect ratio less than 1.35, a circle with half the radius of the original particle was drawn centered around the pixel of maximum intensity. This area was added to a new mask such that only the center of drops was analyzed for area and intensity. The "Analyze particles" tool was used to quantify 'in-drop' areas, and the "Measurement" tool was used to determine background area and total intensity.

Particles were filtered to remove any particle with a Feret diameter (width at widest point) $\leq 4$ pixels and any particle with maximum intensity above 65,500 a.u. (65,535 a.u. is the saturation intensity in 16-bit images; this likely led to an underestimation of dense-phase enzyme and RNA concentrations). For each particle, the average concentration was calculated with the Mean intensity and the calibration curves described above. This analysis was also performed for the background (*i.e.*, dilute phase) and the bulk, non-phase-separated conditions (*i.e.*, no CAPRIN1) (Figure S7B).

Calculating enzyme and RNA concentrations from DVF and PC

The overall average concentrations of fluorescently labeled RNA and enzyme inside and outside drops, generated as described in the preceding section, were then used to calculate the partition coefficient of fluorescently labeled molecules, $PC_{labeled}$, according to

$$PC_{labeled} = \frac{C_{dense,labeled}}{C_{dilute,labeled}} \tag{S35}$$

and droplet volume fraction, $DVF_{labeled}$, or $V_{dense}/V_{total}$, according to

$$DVF_{labeled} = \frac{V_{dense}}{V_{total}} = \frac{C_{total,labeled} - C_{dilute,labeled}}{C_{dense,labeled} - C_{dilute,labeled}} \tag{S36}$$

where $C_{dense,labeled}$ is the concentration of labeled molecule (RNA or enzyme) inside the drop, $C_{dilute,labeled}$ is the concentration of labeled molecule in the background, and $C_{total,labeled}$ is the bulk concentration of fluorescently labeled molecule. Variance from the microscopy measurements was propagated for the error in $PC_{labeled}$ and $DVF_{labeled}$. For a given bulk concentration of CNOT7 (1, 3, or 5 μM) and 100 nM $RNA(A)_{18}$, average values of $PC_{labeled}$ and $DVF_{labeled}$ were calculated, plotted as a function of bulk concentration, and fit with a linear equation. Because some bulk concentrations of enzyme used in the PS-CAP1 assay experiments were not tested via microscopy, the enzyme and RNA $DVF_{labeled}$ and $PC_{labeled}$ were then calculated from the linear fit for each bulk enzyme concentration tested with the FRET assay (Figure S7C).

For the two-phase fits of FRET assay data, it was necessary to calculate the enzyme and RNA concentrations in the dilute ($C_{dilute,assay}$) and dense ($C_{dense,assay}$) phases. To achieve this, the DVF and PC for RNA or enzyme in the FRET assay conditions (*i.e.*, without fluorescently labeled enzyme), $DVF_{assay}$ and $PC_{assay}$, were assumed to be equivalent to the $DVF_{labeled}$ and $PC_{labeled}$ respectively. Therefore, for each bulk enzyme concentration, enzyme and RNA concentrations in the dilute and dense phases were calculated according to

$$C_{dilute,assay} = \frac{C_{total,assay}}{DVF_{labeled}(PC_{labeled}-1)+1} \tag{S37}$$

$$C_{dense,assay} = PC_{labeled} \cdot C_{dilute,assay} \tag{S38}$$

were $C_{total,assay}$ is the total enzyme or RNA concentration used in the assay. See Table S3 for $DVF_{labeled}$, $PC_{labeled}$, $C_{dilute,assay}$, and $C_{dense,assay}$ values used in DEC FRET assays and for fitting PS-CAP1 FRET assay results.

Limitations

Fluorescence microscopy is commonly used to study biomolecular condensates, but dense-phase properties can introduce substantial error in fluorescence measurements by impacting fluorophore photo-physics[14,15]. Fluorophores can be characterized by parameters such as emission wavelength, quantum yield, and fluorescence lifetime, which are sensitive to solvent properties such as crowding, viscosity, pH, and solvent polarity[16,17]. These solvent properties are known to vary within dense phases, which can lead to increased fluorophore quenching[18], decreased fluorescence lifetimes[15,19], and an overall change in measured fluorescence intensity that can result in underestimation of dense-phase protein concentrations when comparing fluorescence in the dense phase to a standard curve generated in dilute conditions[14,17,20] (although concentration overestimation due to fluorophore stabilization should not be ruled out as a possibility depending on condensate properties[16,21]). Furthermore, conjugating organic fluorophores such as Alexa Fluor, cyanine, or BODIPY dyes to proteins can change the phase behavior of the protein[22], meaning a phase-separated system observed via fluorescence may be fundamentally different from a system with the same biomolecules but no fluorophores.

Size of drops can also present an issue for quantitative microscopy. Ideally, one would be able to sample fluorescence intensities from the center of drops where the height is consistent and where one can be confident that the pixels cover a single phase (in the case of multi-phasic condensates).

This would ensure that each pixel sampled covers the same volume of a given phase. In our system, drops were often ~1 μm in diameter, which led to two problems. First, higher concentrations of CAPRIN1, RNA, and CNOT7 lead to multiphasic condensates (data not shown), and these seem to disappear at lower concentrations. However, because the drops in these conditions were so small, it was difficult to say with certainty that each drop was made of a single phase. Rather, the multi-phasic properties may not have been visible with the microscopes available to us. Second, because of the small size, the region of peak intensity only spanned a few pixels. We sampled from the center of drops, but the standard deviation for the fluorescence intensity in each drop was still quite large, suggesting that we were collecting fluorescence data from regions of varying drop height and/or concentration.

For the purposes of this study, which aims to address the conceptual issue of deconvoluting several factors that regulate enzyme activity in condensates, the microscopy and analysis we perform generate reasonable dense- and dilute-phase concentrations of enzyme and RNA. We therefore used these values in our DEC experiments and PS-CAP1 activity analyses.

**Fitting PS-CAP1 deadenylation assay data with dense-phase concentrations**

Dense-phase *vs* dilute-phase deadenylation parameters

To determine the dense-phase $k^*_{obs}$ value, CNOT7 activity in PS-CAP1 conditions was modeled with a two-phase system that includes contributions from both the dilute and dense phases (via eq S3). In this approach, deadenylation reactions in the dense and dilute phases proceeded simultaneously (with eqs S22-S24) based on enzyme and RNA concentrations calculated according to eqs S37 and S38. Note that the same DVF and PC values were used for all poly(A)

RNA species. While fitting, the $k_{obs}^{*}$ value for the dilute phase was set to be the same as that obtained from buffer conditions, and only $k_{obs}^{*}$ for the dense phase was adjusted.

The flux of poly(A) RNA species between dense and dilute regions was calculation over the course of the deadenylation reactions by assuming that the RNA species generated in each phase rapidly equilibrates, reflecting the overall RNA DVF and PC. This is achieved in the two-phase model by first combining the dense- and dilute-phase RNA species, $TA_i$, after each time step to determine the overall concentration of each poly(A) RNA species (this step is also necessary to calculate the FRET). Next, each poly(A) RNA species was then "redistributed" between the phases according to eqs S37 and S38 based on the overall RNA DVF and PC. The newly calculated concentrations of each poly(A) RNA species concentration were then used when solving the rate equations for the next time point.

**Molecular Dynamics**

Generation of initial poses

We created a pool of initial poses of the CNOT7 tail and initial conformations for the CAPRIN1 C-terminal low complexity region, LCR, (CAPRIN1, 103 residues) for molecular dynamics (MD) simulations. IDPConformerGenerator[23,24] was used in conjunction with the experimentally solved X-ray crystal structure of CNOT7 (chain B of PDB 4GMJ[25]) to build the 19-residue C-terminal IDR of the enzyme ("tail", residues 267-SYVQNGTGNAYEEEANKQS-285). The first 9 residues of CNOT7 were not modeled in the crystal structure and, as they don't participate in the dynamics of interest, were left out of our simulations. The reported residue numbering scheme follows that of full length CNOT7 for consistency. IDPConformerGenerator was also used to generate poses of CAPRIN1 used in the simulations.

For the initial “open” pose of CNOT7, the IDR conformations were generated using “ANY” secondary-structure sampling option in IDPConformerGenerator[23] to reflect torsion angle distributions as observed in non-redundant X-ray crystal structures with resolution ≤ 2.0 Å in the RCSB PDB[26] as of November 2024. Generating 10,000 conformers of the CNOT7 tail, we saw an approximately 60% propensity to form α-helical secondary structures within the last 10 residues of the tail (Figure S9A), so we chose a representative helical pose and placed it in the open conformation while defining the open conformation as the tail residues (E278-S285) being at least 12 Å away from $Mg^{2+}$ ions. For the initial “closed” pose, we fixed the positions from the partially modeled tail region as seen in the 4GMJ structure[25] (residues 274-GNAYEEE-280) and modeled the remaining residues (residue ranges 267-273 and 281-285) of the tail with IDPConformerGenerator’s LDRS module[24] with the same secondary structure filter “ANY” as the “open” CNOT7 conformation.

To generate initial poses of the CAPRIN1 LCR, 10,000 conformers were first generated using IDPConformerGenerator using the “ANY” secondary structure filter. The propensity for α-helices was found to be consistently below 35% across the 103-residue long LCR (Figure S9A). For use in the simulations, we chose 7 individual conformations of the CAPRIN1 LCR to be roughly representative of the secondary structure, with largely extended loop conformations.

Simulations

Four separate MD simulations (“open” CNOT7 in buffer and CAPRIN1 condensed phase, and “closed” CNOT7 in buffer and CAPRIN1 condensed phase) were performed using the OpenMM package[27] with the CHARMM36 force field for proteins and the CHARMM36m water model for solvents[28]. For the buffer CNOT7 simulations, the initial protein structure was solvated in a water cube with a side length of 72.7 Å at pH 7.4. The ionic strength of the system was set to 0.147 M with 10 $Cl^-$, 22 $Na^+$, and the existing 2 $Mg^{2+}$ ions. For the CAPRIN1 condensed-phase simulations, the water/solvent cube had a side length of 205 Å, solvating the proteins at pH 7.4. The ionic strength of the system was set to 0.112 M with 331 $Cl^-$, 252 $Na^+$, and the existing 2 $Mg^{2+}$ ions coupled to CNOT7. The 7 CAPRIN1 LCR chains were placed close to CNOT7 in the starting conformation, with the first part of the simulation leading to greater coalescence around CNOT7, effectively simulating a 30 mM dense-phase environment[29].

For both "open" and "closed" CNOT7 in the absence and presence of CAPRIN1, energy minimization was performed, after which the systems were gradually heated from 50 K to 300 K in 50 K increments. Subsequently, the systems were equilibrated for 20 ns under the NVT ensemble (constant number of particles, volume, and temperature), followed by an additional 20 ns under the NPT ensemble (constant number of particles, pressure, and temperature). The production simulations were then carried out for 1 μs for the CNOT7 with CAPRIN1 chains and 2 μs for CNOT7 in the buffer condition, for both "open" and "closed" initial poses (Figure 5A and 5B).

Simulation analysis

Analyses of the simulation trajectories were performed using an in-house Python script (provided in the supplemental archive) using the MDTraj[30] library. Residue-residue and residue-ion distances as well as position in space were calculated by using the "compute center of mass" function within MDTraj. Interaction times were defined as the total time that the centers of mass of particular CAPRIN1 and CNOT7 residues (or ions) were ≤ 6 Å from each other (Figures 5C and S10).

CNOT7 diffusion was quantified by applying MDTraj "compute center of mass" to the full CNOT7 protein in each simulation to obtain 3D coordinates for tracking position over time (Figure S11A). Positions were corrected to remove any frame-to-frame (1 ns) jumps > 5 Å. Mean squared displacement (MSD, $Å^2$) of the center of mass was calculated according to

$$MSD(\Delta t) = \langle (r(t+\Delta t) - r(t))^2 \rangle_{0 \le t < t_{total}-\Delta t}, 0 \le \Delta t < t_{total}/2 \quad \text{(S39)}$$

where $\Delta t$ is the lag time (ns) and $t_{total}$ is the total duration of the simulation (1000 ns, Figure S11B). The MSD was trimmed to 200 ns lag time and fit with the linear equation

$$MSD(\Delta t) = 2nD\Delta t \quad \text{(S40)}$$

where $n$ is the number of dimensions (in this case $n$ = 3) and $D$ is the diffusion coefficient in $Å^2$/ns (Figure 5D, Figure S11B)[31].

The final pose from the buffer (open start position) simulation was aligned to the AlphaFold2 structure of CNOT7 (AF-Q9UIV1-F1-v6)[32] using PyMOL[33] via residues 29-AMDTEFPG-36, 145-LSFHSGYDFG-154, and 214-PQHQAGSDSL-223 (Figure S9B). These sequences include residues 31-34, 146-148, 151-152, 216-218, and 221, which are within 5 Å of the active site $Mg^{2+}$ ions. The additional residues were included to accommodate PyMOL's alignment protocol.

**Evolutionary signature analysis**

Evolutionary conservation of CAPRIN1 LCR bulk features was determined by calculating evolutionary signatures using what is now called FAIDR-ES[34] for the full CAPRIN1 IDR2 (247-709) and for the last 100 residues to approximate the LCR. CAPRIN1 orthologues for all species were downloaded from the Ensemble REST server[26] and filtered using previously reported methods[27]. Evolutionary signatures were quantified by Z-scores (Table S5), which indicate enrichment/more prevalent in/larger (+ values) or depletion/less prevalent in/smaller (- values) of a given feature in an IDR of a protein and its orthologs compared to a distribution of random computationally evolved sequences (without conservation of features).

**Other computational tools**

Images of protein structures were generated with PyMOL (Figures 5A, 5B, and S9B). Data plots were generated using the *Matplotlib* Python library[35]. Deadenylation assay fits and simulations were carried out using the *deadenylationkinetics* software package (github.com/rosieirwin)[8]. Unpaired *t* tests were performed using *SciPy* Python library[10]. Protein Calculator[36] was used to determine protein net charge at pH 7.4.

### SUPPLEMENTARY DATA

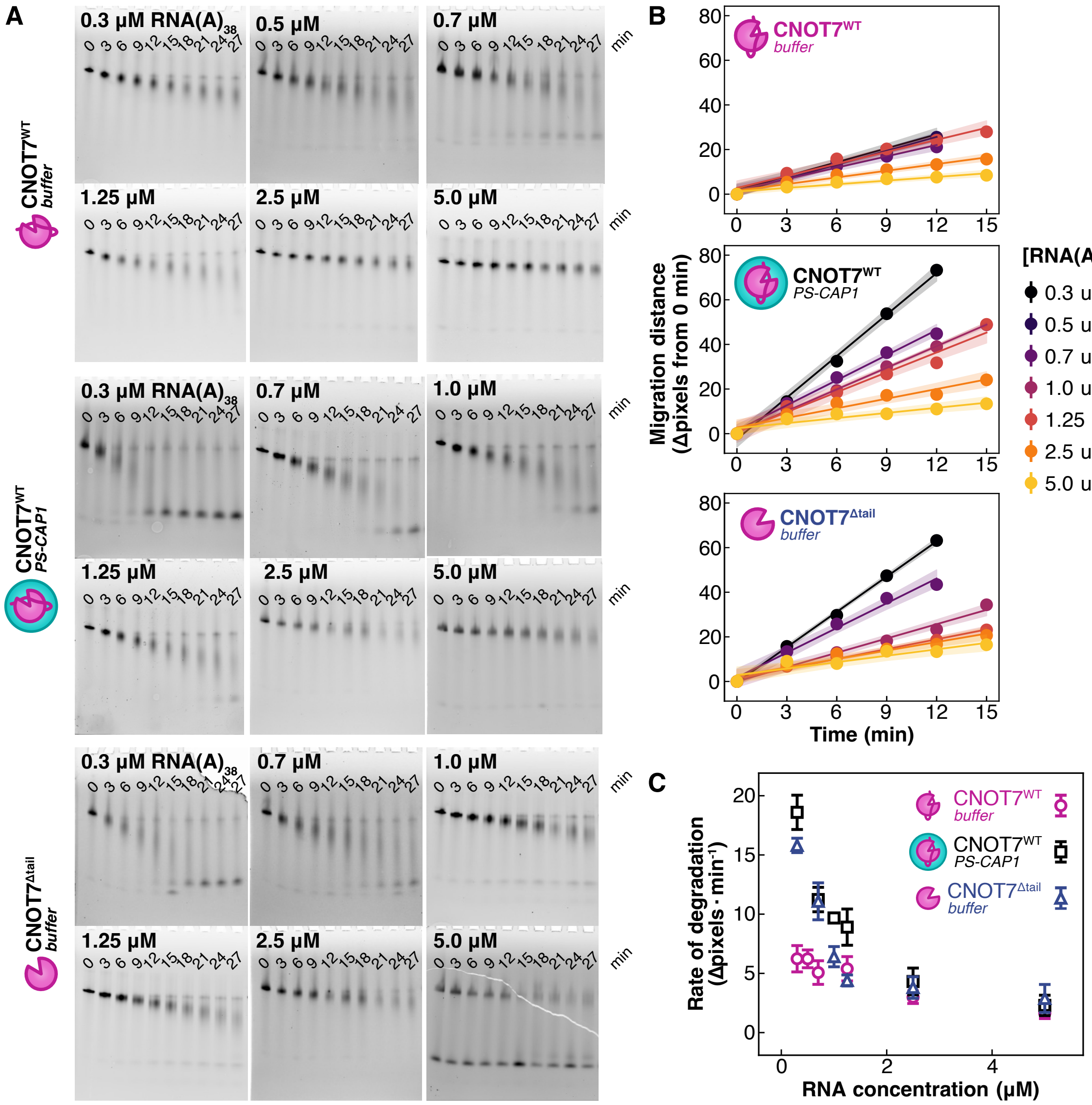


**Figure S1.** CNOT7 activity is modulated by its tail and phase separation with CAPRIN1. (A) Denaturing gels imaged to detect FITC fluorescence from the 5' FITC-CCUUUCC(A)$_{38}$ RNA substrate (RNA(A)$_{38}$, 0.3-5 μM). The bulk enzyme and CAPRIN1 concentrations were 1 μM and 100 μM respectively in all conditions. Images were analyzed with ImageJ to quantify migration of the bands (corresponding RNA with decreasing poly(A) tail length) with respect to the band intensity at t = 0 min (in terms of number of pixels traversed). (B) Migration distances from A as a function of time, with linear fits of the data. The shaded areas surrounding each fitted line correspond to the 95% confidence intervals. (C) Profiles of RNA(A)$_{38}$ degradation rates (pixels/min) *vs* [RNA] from Figure S1B. The rates, in terms of pixels traversed, converge when [RNA$(A)_{38}$] > 2 μM because more time is required for the enzyme to consume high quantities of full-length RNA. Error bars show 95% confidence intervals (CI) from linear fits of gel data from B.

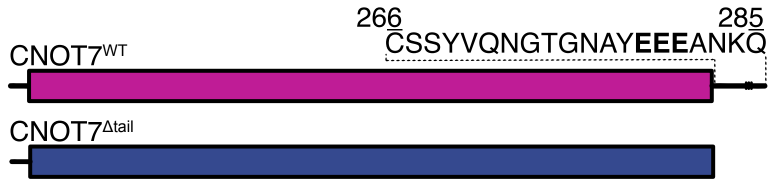


**Figure S2.** CNOT7 constructs. Constructs include CNOT7$^{WT}$ (1-285) and CNOT7$^{\Delta tail}$ (1-265).

**A**

| Constant | Value |
|---|---|
| $k_1$ | $1\times10^{-2}$ $s^{-1}$ (B), 100 $s^{-1}$ (C) |
| $k_{-1}$ | $1\times10^{-2}$ $s^{-1}$ (B), 100 $s^{-1}$ (C) |
| $k_2$ | $1\times10^{5}$ $M^{-1}s^{-1}$ |
| $k_{-2}$ | 3 $s^{-1}$ |
| $k_{cat}$ | $1.15\times10^{-1}$ $s^{-1}$ |
| $k^*_{obs}$ | $1.85\times10^{3}$ $M^{-1}s^{-1}$ |

$$K^* = \frac{k_{-1}}{k_1}$$

$$K_M = \frac{k_{-2} + k_{cat}}{k_2}$$

$$k^*_{obs} = \frac{k_{cat}}{K_M(1 + K^*)}$$

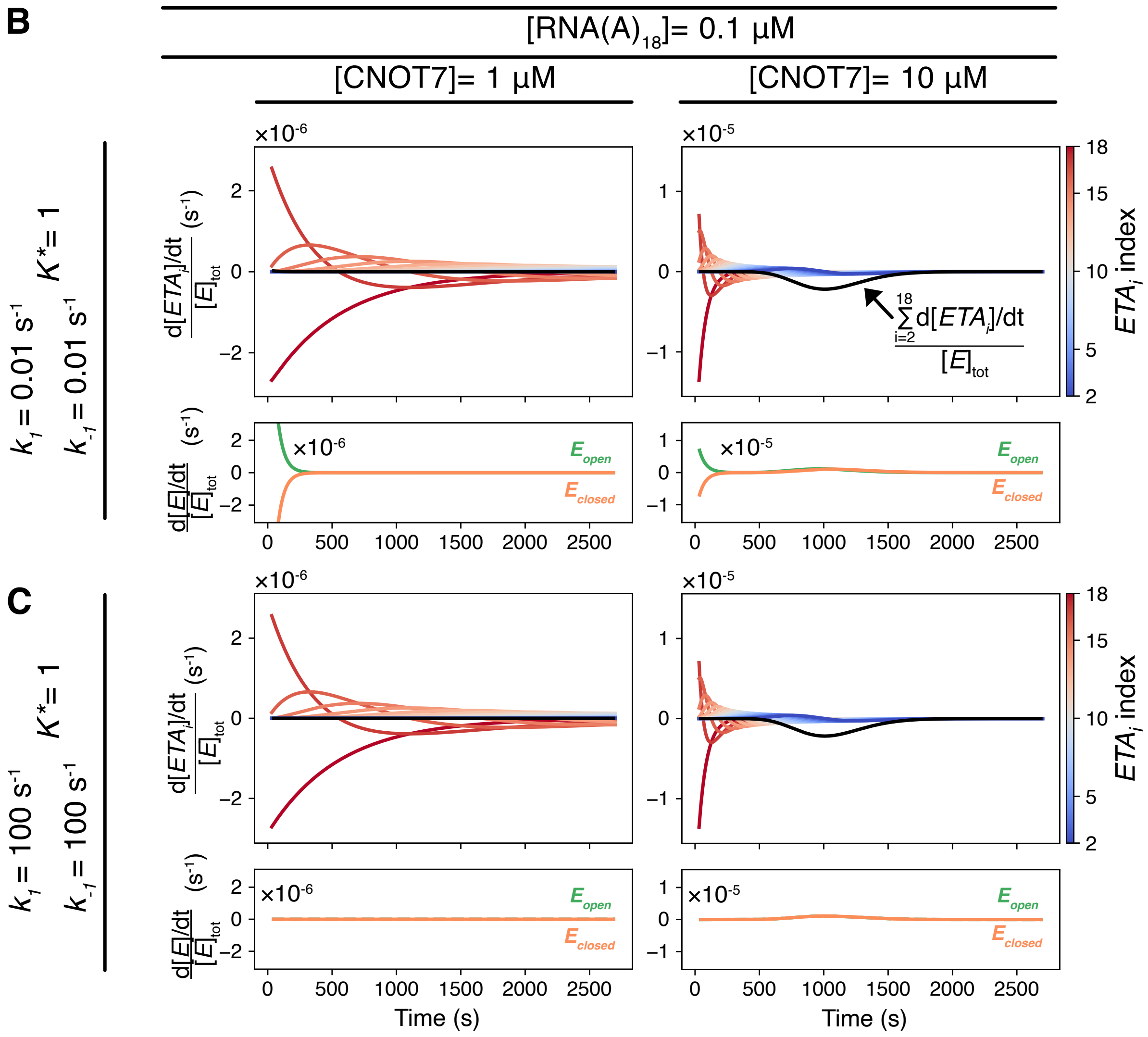


**Figure S3.** Validation of the steady state assumption used in the derivation of the kinetic deadenylation model. (A) Table of rate constants used to simulate the data in (B-C). The calculation of $k^*_{obs}$ using the constants from the table is shown to the right. Reaction rates for the free and RNA-bound enzyme species were simulated with slow tail dynamics, that is $k_1$ and $k_{-1}$ both set to 0.01 $s^{-1}$ along the top row (B) at an initial RNA$(A)_{18}$ concentration of 0.1 μM and a concentration for CNOT7 of either 1 μM (left) or 10 μM (right). The reaction rates were normalized by the enzyme concentration to show that their magnitudes are small and roughly constant relative to the total amount of enzyme, confirming the steady state assumption. Identical reactions were simulated along the bottom row (C) except with rapid tail dynamics corresponding to 100 $s^{-1}$ for $k_1$ and $k_{-1}$. The black lines are the sum of the rates for the RNA-bound enzyme species that are detected in the FRET assay. Curves for longest to shortest bound RNAs are shown in red to blue. The time courses for $E_{closed}$ and $E_{open}$ are shown in the subpanels as orange and green lines respectively. The $k_1$ and $k_{-1}$ were both set to either 0.01 or 100 $s^{-1}$, so as to maintain a $K^*$ value of 1, corresponding to 50% closed CNOT7 at equilibrium, to explore the influence of the tail dynamics on the reaction curves and establish how $[E_{closed}]$ and $[E_{open}]$ vary throughout the course of each of the trajectories. Values of $k_2$, $k_{-2}$, and $k_{cat}$ that govern binding of RNA, dissociation of RNA-enzyme complexes, and RNA cleavage respectively were selected to produce simulated FRET profiles that were similar to the experimental progress curves. The full set of 5 rate constants was ultimately constrained so that calculated $k^*_{obs}$ values were similar to experimentally determined $k^*_{obs}$ for CNOT7 at 25 °C in buffer ($1.82\times10^3$ $M^{-1}$ $s^{-1}$).

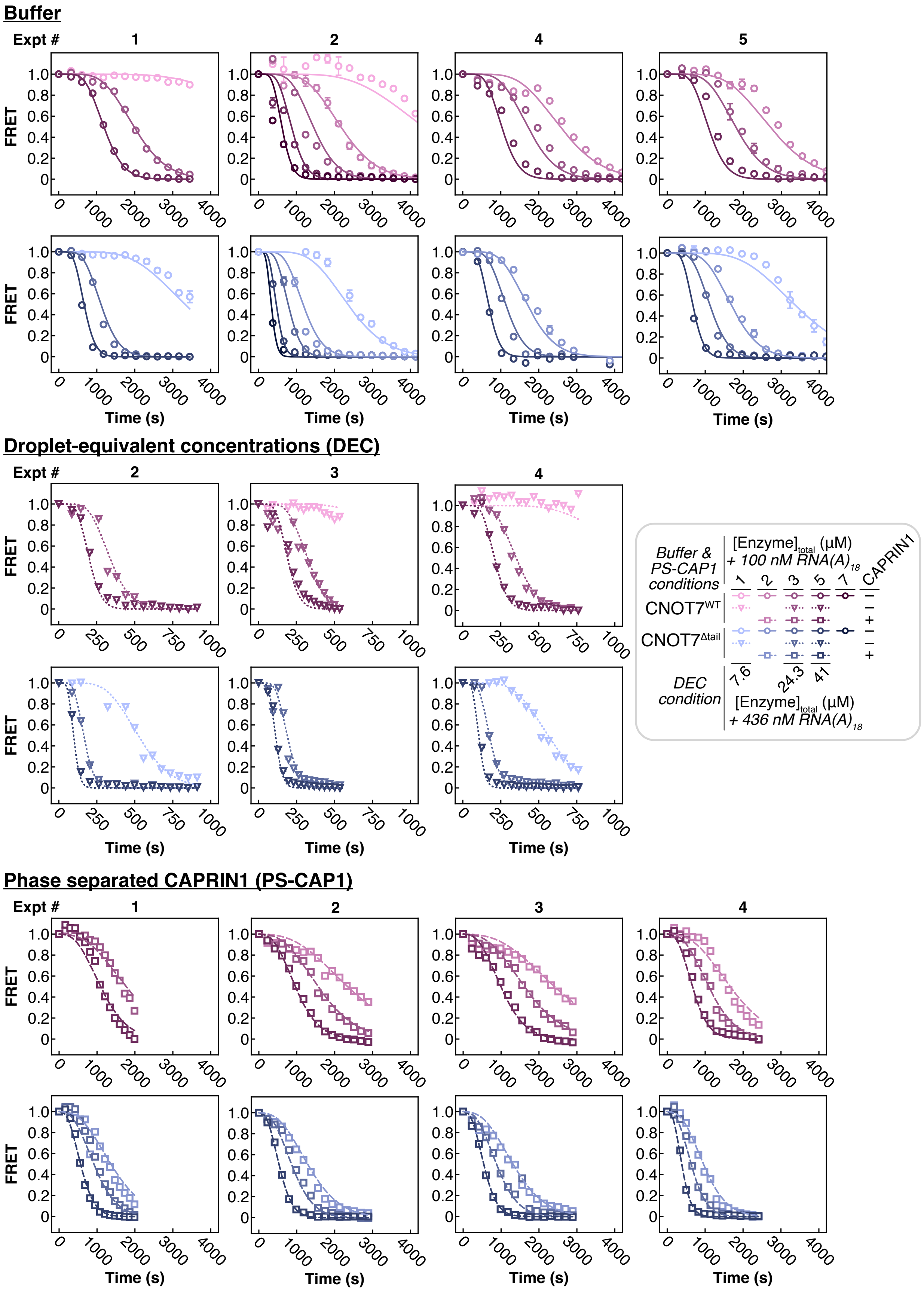
Buffer
Expt #
1
2
4
5
FRET
Time (s)
Droplet-equivalent concentrations (DEC)
Expt #
2
3
4
FRET
Time (s)
Buffer &
PS-CAP1
conditions
[Enzyme]total (μM)
+ 100 nM RNA(A)18
1
2
3
5
7
CAPRIN1
CNOT7WT
CNOT7Δtail
DEC
condition
7.6
24.3
41
[Enzyme]total (μM)
+ 436 nM RNA(A)18
Phase separated CAPRIN1 (PS-CAP1)
Expt #
1
2
3
4
FRET
Time (s)

**Figure S4.** All data for degradation of RNA(A)$_{18}$ by CNOT7$^{WT}$ (pink) or CNOT7$^{\Delta tail}$ (blue) in buffer (circles, solid lines), at the CAPRIN1 droplet-equivalent concentration (DEC, inverted triangles, dotted lines), or PS-CAP1 (squares, dashed lines) at 25 °C. Experiment numbers differentiate each replicate; buffer experiment 3, PS-CAP1 experiment 5, and DEC experiment 1 are shown in Figure 4A. Note that the time for reaction completion in DEC conditions is much shorter in general than in buffer or PS-CAP1 (see time axes).

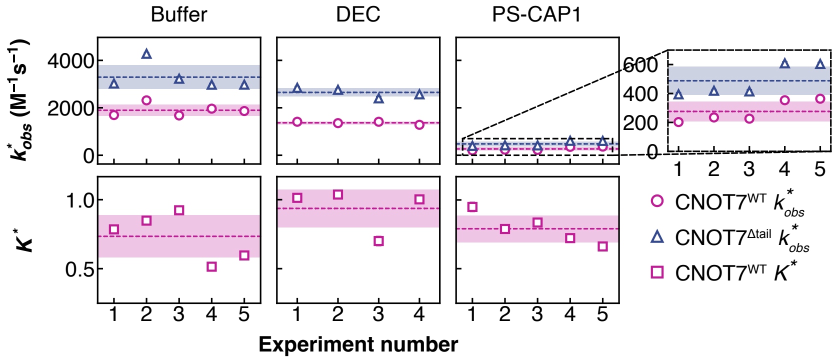


**Figure S5.** Fit results for all assay data. $k^*_{obs}$ values derived from fits of all deadenylation experiments with our kinetic model for CNOT7$^{WT}$ (pink, circles) and CNOT7$^{\Delta tail}$ (blue, triangles) in buffer, DEC, and PS-CAP1 conditions. $K^*$ of CNOT7$^{WT}$ (pink, squares) were calculated from CNOT7$^{WT}$ and CNOT7$^{\Delta tail}$ for each experiment (eq S26). Dashed lines show average values and shaded regions show standard deviation for CNOT7$^{WT}$ (pink) and CNOT7$^{\Delta tail}$ (blue). Experiment numbers correspond to the experiment numbers above the FRET curves in Figure S4, except for buffer experiment 3, PS-CAP1 experiment 5, and DEC experiment 1, which are shown in Figure 4A.

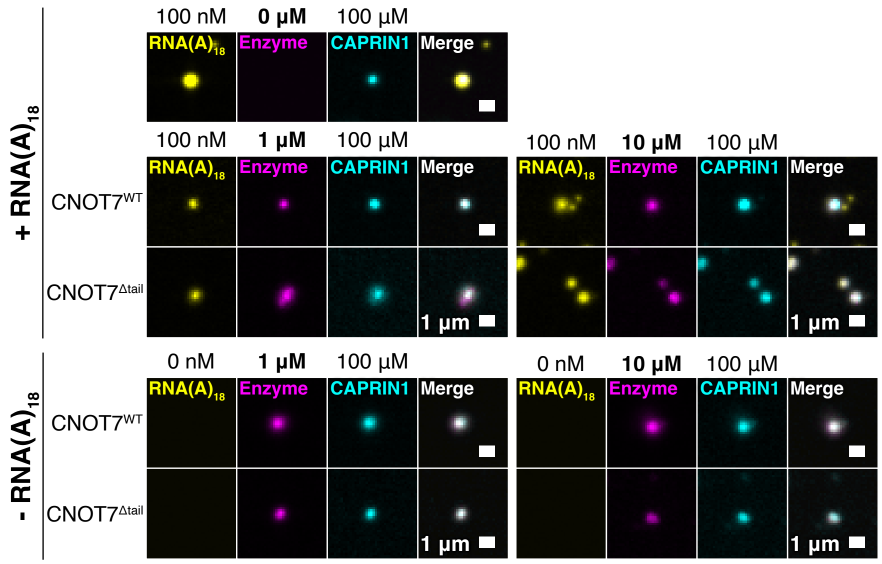


**Figure S6.** CNOT7$^{WT}$ and CNOT7$^{\Delta tail}$ partition with RNA(A)$_{18}$ in CAPRIN1 condensates. Representative confocal microscopy images of condensates with AF647-CAPRIN1 (100 µM, cyan) ± 6FAM-RNA(A)$_{18}$ (100 nM, yellow) ± AF568-enzyme (1 or 10 µM, magenta). Images were acquired on a Leica DMi8 spinning disk confocal microscope equipped with a 63×/1.3 NA oil immersion objective at ambient temperature. Samples were placed in a 96-well black cell imaging plate with glass bottom. Scale bar is 1 µm.

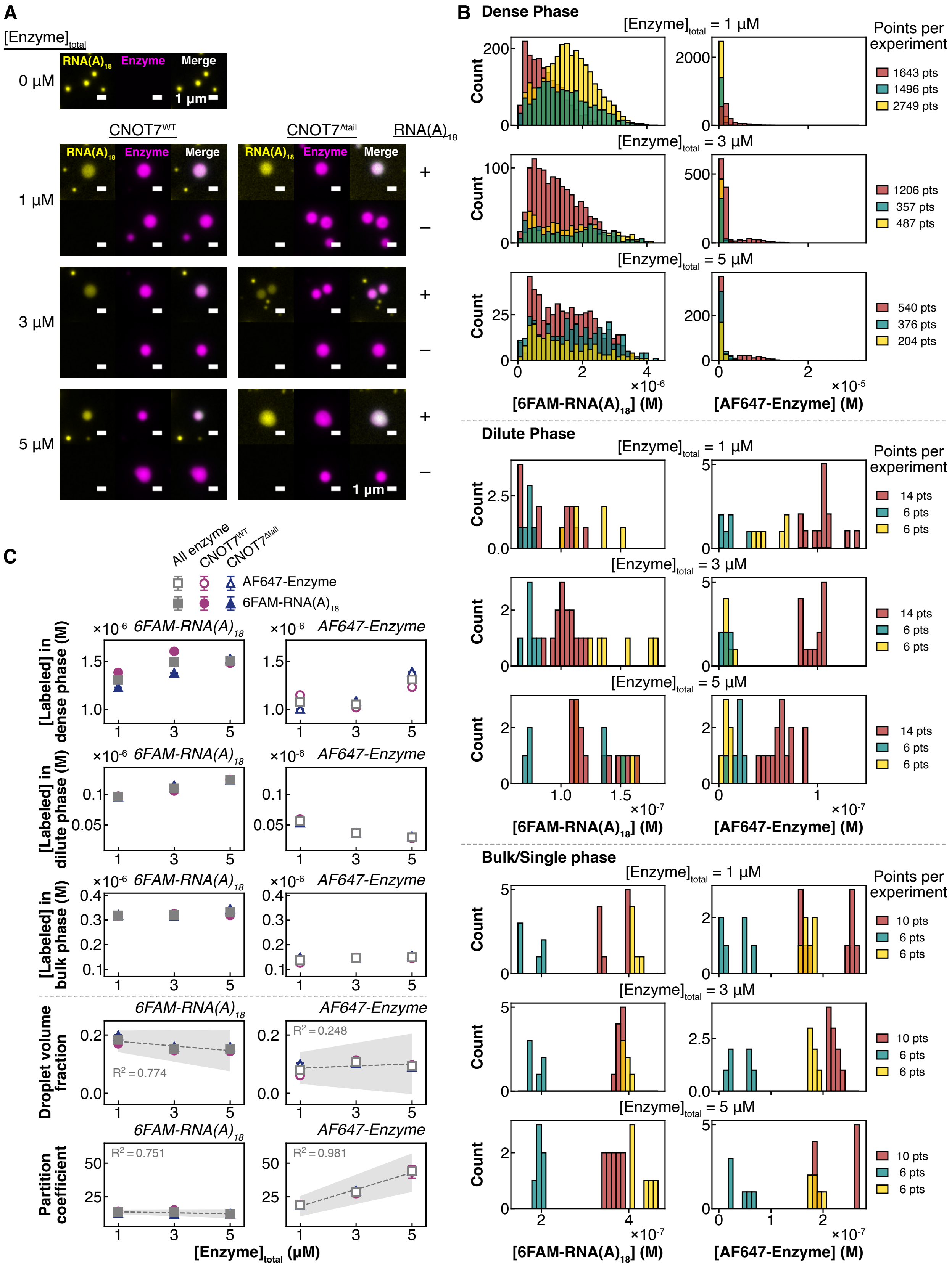
A
[Enzyme]total
0 µM
RNA(A)18
Enzyme
Merge
1 µm
CNOT7WT
CNOT7Δtail
RNA(A)18
1 µM
3 µM
5 µM
+
–
B
Dense Phase
[Enzyme]total = 1 µM
[Enzyme]total = 3 µM
[Enzyme]total = 5 µM
Count
Points per experiment
1643 pts
1496 pts
2749 pts
1206 pts
357 pts
487 pts
540 pts
376 pts
204 pts
×10-6
×10-5
[6FAM-RNA(A)18] (M)
[AF647-Enzyme] (M)
Dilute Phase
14 pts
6 pts
6 pts
×10-7
Bulk/Single phase
10 pts
6 pts
6 pts
C
All enzyme
CNOT7WT
CNOT7Δtail
AF647-Enzyme
6FAM-RNA(A)18
[Labeled] in dense phase (M)
[Labeled] in dilute phase (M)
[Labeled] in bulk phase (M)
Droplet volume fraction
Partition coefficient
R2 = 0.774
R2 = 0.248
R2 = 0.751
R2 = 0.981
[Enzyme]total (µM)

**Figure S7.** Fluorescence microscopy allows for determination of droplet volume fraction (DVF) and partition coefficients (PCs) for CNOT7 and $RNA(A)_{18}$ in phase-separated CAPRIN1 conditions. (A) Representative confocal microscopy images for enzyme-RNA colocalization. Sample contained 6FAM-$RNA(A)_{18}$ (100 nM), unlabeled CAPRIN1 (100 µM), and $CNOT7^{WT}$ or $CNOT7^{\Delta tail}$ (1, 3, or 5 µM with 50 nM AF647-labeled enzyme) and were imaged at ambient temperature in a PF127-passivated 96-well glass bottom plate on a Zeiss Elyra PS.1 microscope (Zeiss) equipped with a 100×/1.46 NA oil immersion objective. All condensates imaged were settled on the glass well-bottom. Dilute-phase and bulk-solution images (images not shown) were acquired several µm above the bottom of the well. (B) Quantification of AF647-enzyme and 6FAM-$RNA(A)_{18}$ dilute-phase, dense-phase, and bulk concentrations. Images were analyzed using an in-house ImageJ script. Background-corrected intensities were then converted to concentrations with a standard curve. Histograms show concentration values determined for each drop, dilute phase, or bulk (single) phase imaged. Experiments were performed in triplicate (red, teal, yellow), and for each experiment, data for conditions with similar enzyme concentrations (*e.g.*, 1 µM $CNOT7^{WT}$ and 1 µM $CNOT7^{\Delta tail}$) were combined. (C) Average dense-phase, dilute-phase, and bulk concentrations for fluorescently labeled enzyme (empty markers) or $RNA(A)_{18}$ (solid markers). $CNOT7^{WT}$ (pink circles) and $CNOT7^{\Delta tail}$ (blue triangles) were initially analyzed separately, but due to insignificant differences in concentration values, the datasets for each condition were combined to determine the overall average concentrations (grey squares) of labeled enzyme or labeled $RNA(A)_{18}$. These values were used to determine the partition coefficients and droplet-volume fractions for each total enzyme concentration. The grey dashed line shows a linear regression of the average points, and the shaded region indicates the standard error of the fit. The DVFs and PCs for unlabeled enzyme and RNA were assumed to be the same as the DVF and PC for fluorescently labeled enzyme and RNA determined here.

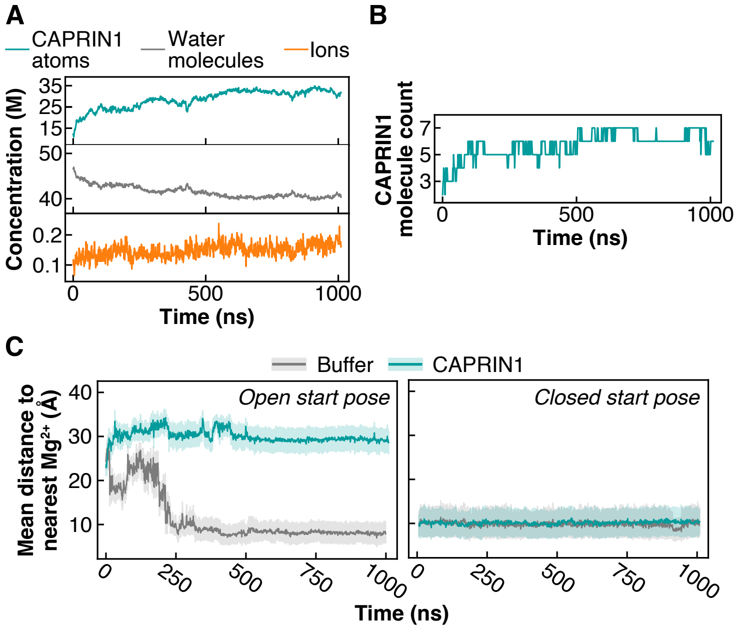


**Figure S8.** Analysis of molecular dynamics trajectories for CNOT7$^{WT}$ in buffer or CAPRIN1. (A) Concentration of water molecules (grey), CAPRIN1 atoms (teal), and charged ions (orange) over 1 µs of MD simulation within a box with edge length = 72.7 Å centered around CNOT7. (B) CAPRIN1 chain count within the box over 1 µs. A chain was counted if its center of mass fell within the box. (C) Average distances of CNOT7 tail residues E278, E279, and E280 from the closest active site $Mg^{2+}$ ion in buffer conditions (grey) or with CAPRIN1 (teal) when simulations were initiated with CNOT7 in an open (left) or closed (right) conformation for 1 µs. Shaded regions indicate standard deviation.

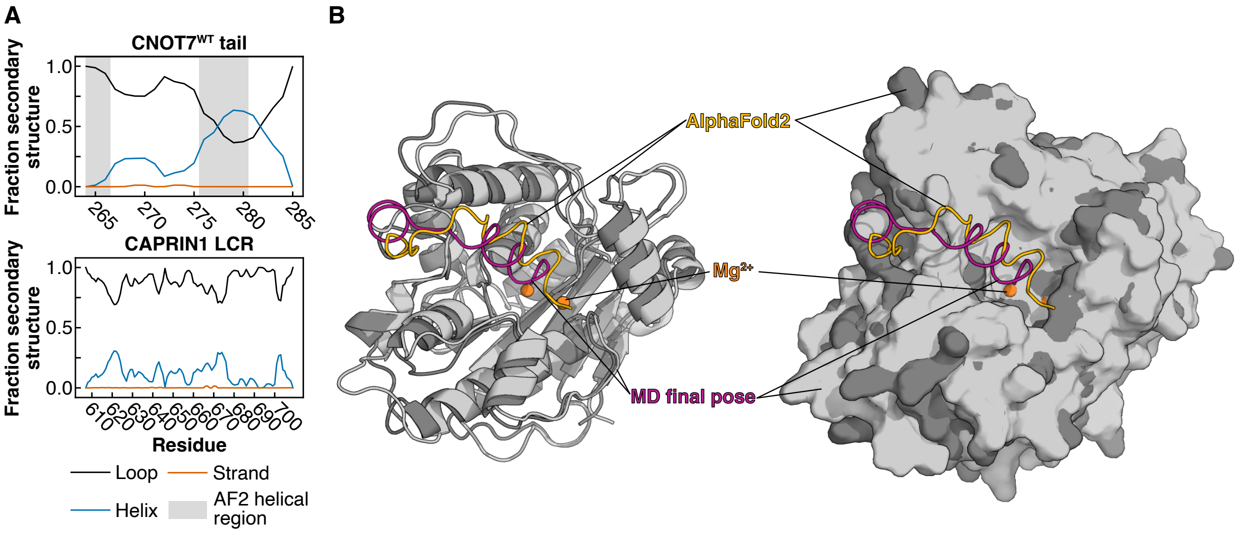


**Figure S9.** Structural similarities between MD/IDPConformerGenerator CNOT7 and AlphaFold2 CNOT7. (A) Secondary structure propensity of the CNOT7 tail or CAPRIN1 LCR. 10,000 conformers of each protein were generated with IDPConformerGenerator, with the CNOT7 tail built in the context of the folded domain. Grey shaded regions indicate helical propensity for the tail of the AlphaFold2 structure. DSSP[37] secondary structure annotation (based on hydrogen-bonding patterns) is plotted for Helix (blue, combined codes: "H" alpha helix, "G" 310 helix), Strand (orange, "E" beta strand), and Loop (black, combined codes: "_" loop or coil, "?" loop or coil, "P" PPII helix, "T" H-bonded turn, "S" bend, "B" beta bridge, "I" pi helix). (B) Ribbon and surface alignments of the final pose of CNOT7 from the buffer simulation (pink, light grey) and the AlphaFold2 CNOT7 structure (AF-Q9UIV1-F1-v6; yellow, dark grey). Structures were aligned via residues 29-36, 145-154, and 214-223, which are near the $Mg^{2+}$ ions (orange). Yellow region of AF2 structure has a pLDDT score < 50.

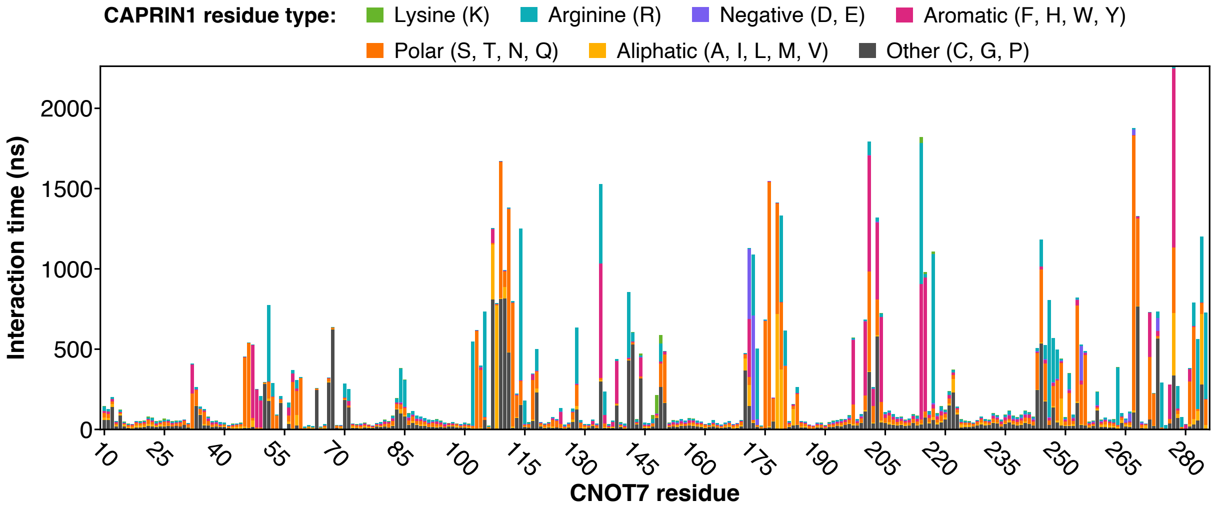


**Figure S10.** Interaction times for individual CNOT7 residues with CAPRIN1 residues by category. Residues were considered to be interacting if their centers of masse were separated by ≤ 6 Å for ≥ 5 ns. Residues 1-9 were excluded from the simulation.

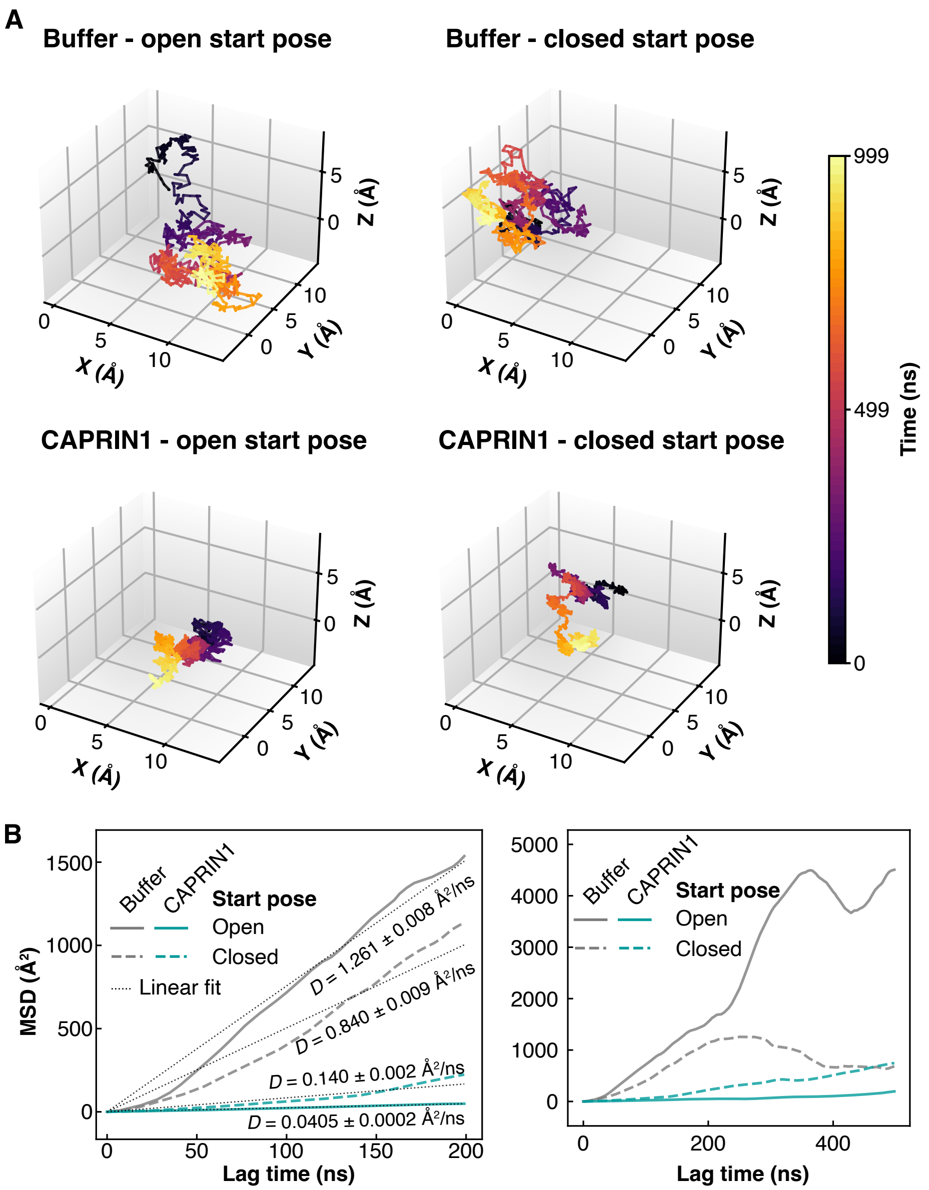


**Figure S11.** CNOT7 diffusion during MD simulations. (A) Position of CNOT7 center of mass for CNOT7 in buffer or CAPRIN1 with open or closed start poses. Color of lines corresponds to time (ns) shown in the color bar. (B) Mean squared displacement (Å$^2$) of CNOT7 center of mass between simulation time $t$ and time $t + \Delta t$ where $\Delta t$ is the lag time (ns) for CNOT7 in buffer (grey) or CAPRIN1 (teal) conditions with an open (solid) or closed (dashed) start pose. $R^2$ values for fits are: buffer (open start pose) 0.976, buffer (closed start pose) 0.938, CAPRIN1 (open start pose) 0.985, CAPRIN1 (closed start pose) 0.872. Full MSD dataset (right) was cut at 200 ns lag time for linear fits (dashed lines, left).

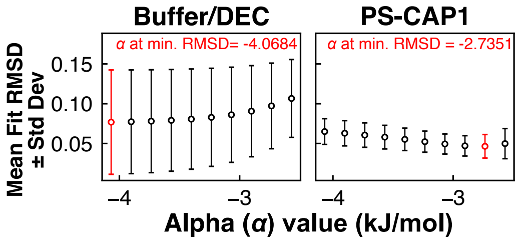


**Figure S12.** Optimization of $\alpha$ RNA-DNA hybridization parameter for buffer and PS-CAP1 conditions (eq S29). Buffer and PS-CAP1 FRET assay data for CNOT7$^{WT}$ and CNOT7$^{\Delta tail}$ were fit with an array of $\alpha$ (alpha) values, and the average RMSDs for all CNOT7$^{WT}$ and CNOT7$^{\Delta tail}$ fits in each condition were determined. The $\alpha$ with the lowest RMSD, indicated in red, was used for the final fits.

**Table S1.** Extracted $k^*_{obs}$, $K^*$, $k_{cat}/K_M$ values from fits of time dependent FRET recorded under buffer, PS-CAP1, and DEC conditions.

| Enzyme | [Enzyme]$_{total}$ (µM) | Condition | $k^*_{obs}$ ± Std. Dev. (M$^{-1}$s$^{-1}$) | $K^*$ ± Std. Dev |
|---|---|---|---|---|
| CNOT7$^{WT}$ | 1-7 | buffer | 1900 ± 200 | 0.7 ± 0.2 |
| CNOT7$^{\Delta tail}$ | 1-7 | buffer | 3300 ± 500 | 0 |
| CNOT7$^{WT}$ | 1-5 | PS-CAP1 | 280 ± 70 | 0.8 ± 0.1 |
| CNOT7$^{\Delta tail}$ | 1-5 | PS-CAP1 | 490 ± 90 | 0 |
| CNOT7$^{WT}$ | 7.6-41 | DEC | 1360 ± 50 | 0.9 ± 0.1 |
| CNOT7$^{\Delta tail}$ | 7.6-41 | DEC | 2600 ± 200 | 0 |

See Table S3 for dense-phase/DEC enzyme and RNA concentrations; [Enzyme]$_{total}$ refers to the bulk enzyme concentration before phase separation. PS-CAP1 = phase-separated CAPRIN1 conditions, DEC = Droplet-equivalent concentration conditions (in the absence of CAPRIN1). $K^*$ values for CNOT7$^{\Delta tail}$ are set to 0 as only the open state can form. As a result, the $k_{cat}/K_M$ values for CNOT7$^{WT}$ are equivalent to the corresponding $k^*_{obs}$ values for CNOT7$^{\Delta tail}$ (see description of the kinetic model above).

**Table S2.** Enzyme and RNA droplet volume fractions (DVF) and partition coefficients (PC) used for fitting PS-CAP1 data.

| $[\text{Enzyme}]_{\text{total}}$ (µM) | Enzyme DVF | Enzyme PC | RNA DVF | RNA PC |
|---|---|---|---|---|
| 1 | 0.08 ± 0.03 | 18 ± 3 | 0.18 ± 0.02 | 13.8 ± 0.9 |
| 2 | 0.09 ± 0.03 | 25 ± 5 | 0.17 ± 0.02 | 13 ± 1 |
| 3 | 0.09 ± 0.04 | 30 ± 6 | 0.16 ± 0.03 | 13 ± 1 |
| 5 | 0.10 ± 0.05 | 43 ± 7 | 0.15 ± 0.04 | 12 ± 2 |

Values were calculated from linear fits in Figure S7C.

**Table S3.** Dilute- and dense-phase enzyme and RNA concentrations used for DEC experiments and fitting PS-CAP1 FRET assay data with a two-phase kinetic model.

| $[\text{Enzyme}]_{\text{total}}$ (µM) | $[\text{Enzyme}]_{\text{dense}}$ (µM) | $[\text{RNA(A)}_{18}]_{\text{dense}}$ (nM) | $[\text{Enzyme}]_{\text{dilute}}$ (µM) | $[\text{RNA(A)}_{18}]_{\text{dilute}}$ (nM) |
|---|---|---|---|---|
| **1** | 7.6 | 421 | 0.405 | 30.4 |
| **2** | 15.7 | 431 | 0.646 | 32.0 |
| **3** | 24.3 | 442 | 0.797 | 33.6 |
| **5** | 41.0 | 465 | 0.955 | 37.4 |

Values calculated with eqs S37 and S38 using values in Table S2. Because RNA concentrations for all total enzyme concentrations were similar, the average dense-phase RNA concentration of 436 nM was used for DEC experiments.

**Table S4.** Separation of contributions to changes in $k^{*}_{obs}$ from changes in tail equilibria and $k_{cat}/K_M$ between conditions *i* and *j*.

| Conditions Compared (*i*/*j*) | $\beta$ ± SEM | $\gamma$ ± SEM |
|---|---|---|
| buffer/buffer | 1.0 | 1 |
| DEC/buffer | 1.1 ± 0.1 | 0.6 ± 0.1 |
| PS-CAP1/buffer | 1.1 ± 0.1 | 0.14 ± 0.04 |
| PS-CAP1/DEC | | |

Shown are $\beta = (1 + K^{*}_{j})/(1 + K^{*}_{i})$ and $\gamma = (k_{cat}/K_M)_i/(k_{cat}/K_M)_j$. See *'Evaluating changes in catalytic efficiency'* above for further details. SEM = standard error of the mean. Data plotted in Figure 4E.

**Table S5.** (separate file) Evolutionary signatures of CAPRIN1 IDR2 (247-709) and LCR (607-709). Each feature is attributed a mean Z-score. Positive (negative) Z-scores indicate a feature is more (less) prevalent in or larger (smaller) for CAPRIN1 homologs than would be expected for random evolution of an intrinsically disordered protein region.

(3) Sattler, M.; Schleucher, J.; Griesinger, C. Heteronuclear Multidimensional NMR Experiments for the Structure Determination of Proteins in Solution Employing Pulsed Field

Gradients. *Prog. Nucl. Magn. Reson. Spectrosc.* **1999**, *34* (2), 93–158. https://doi.org/10.1016/s0079-6565(98)00025-9.

(10) Virtanen, P.; Gommers, R.; Oliphant, T. E.; Haberland, M.; Reddy, T.; Cournapeau, D.; Burovski, E.; Peterson, P.; Weckesser, W.; Bright, J.; Walt, S. J. van der; Brett, M.; Wilson, J.; Millman, K. J.; Mayorov, N.; Nelson, A. R. J.; Jones, E.; Kern, R.; Larson, E.; Carey, C. J.; Polat, İ.; Feng, Y.; Moore, E. W.; VanderPlas, J.; Laxalde, D.; Perktold, J.; Cimrman, R.; Henriksen, I.; Quintero, E. A.; Harris, C. R.; Archibald, A. M.; Ribeiro, A. H.; Pedregosa, F.; Mulbregt, P. van; Contributors, S. 1 0; Vijaykumar, A.; Bardelli, A. P.; Rothberg, A.; Hilboll, A.; Kloeckner, A.; Scopatz, A.; Lee, A.; Rokem, A.; Woods, C. N.; Fulton, C.; Masson, C.; Häggström, C.; Fitzgerald, C.; Nicholson, D. A.; Hagen, D. R.; Pasechnik, D. V.; Olivetti, E.; Martin, E.; Wieser, E.; Silva, F.; Lenders, F.; Wilhelm, F.; Young, G.; Price, G. A.; Ingold, G.-L.; Allen, G. E.; Lee, G. R.; Audren, H.; Probst, I.; Dietrich, J. P.; Silterra, J.; Webber, J. T.; Slavič, J.; Nothman, J.; Buchner, J.; Kulick, J.; Schönberger, J. L.; Cardoso, J. V. de M.; Reimer, J.; Harrington, J.; Rodríguez, J. L. C.; Nunez-Iglesias, J.; Kuczynski, J.; Tritz, K.; Thoma, M.; Newville, M.; Kümmerer, M.; Bolingbroke, M.; Tartre, M.; Pak, M.; Smith, N. J.; Nowaczyk, N.; Shebanov, N.; Pavlyk, O.; Brodtkorb, P. A.; Lee, P.; McGibbon, R. T.; Feldbauer, R.; Lewis, S.; Tygier, S.; Sievert, S.; Vigna, S.; Peterson, S.; More, S.; Pudlik, T.; Oshima, T.; Pingel, T. J.; Robitaille, T. P.; Spura, T.; Jones, T. R.; Cera, T.; Leslie, T.; Zito, T.; Krauss, T.; Upadhyay, U.; Halchenko, Y. O.; Vázquez-Baeza, Y. SciPy 1.0: Fundamental Algorithms for Scientific Computing in Python. *Nat. Methods* **2020**, *17* (3), 261–272. https://doi.org/10.1038/s41592-019-0686-2.

(11) Yao, R.-W.; Rosen, M. K. Advanced Surface Passivation for High-Sensitivity Studies of Biomolecular Condensates. *Proc. Natl. Acad. Sci.* **2024**, *121* (22), e2403013121. https://doi.org/10.1073/pnas.2403013121.

(12) Schindelin, J.; Arganda-Carreras, I.; Frise, E.; Kaynig, V.; Longair, M.; Pietzsch, T.; Preibisch, S.; Rueden, C.; Saalfeld, S.; Schmid, B.; Tinevez, J.-Y.; White, D. J.; Hartenstein, V.; Eliceiri, K.; Tomancak, P.; Cardona, A. Fiji: An Open-Source Platform for Biological-Image Analysis. *Nat. Methods* **2012**, *9* (7), 676–682. https://doi.org/10.1038/nmeth.2019.

(13) Schmid, M. *Adjustable Watershed*; 2022.

(24) Liu, Z. H.; Teixeira, J. M. C.; Zhang, O.; Tsangaris, T. E.; Li, J.; Gradinaru, C. C.; Head-Gordon, T.; Forman-Kay, J. D. Local Disordered Region Sampling (LDRS) for Ensemble

Modeling of Proteins with Experimentally Undetermined or Low Confidence Prediction Segments. *Bioinformatics* **2023**, *39* (12), btad739. https://doi.org/10.1093/bioinformatics/btad739.

(33) Schrödinger, L. The PyMOL Molecular Graphics System, Version~3.1. https://www.pymol.org/.

(34) Zarin, T.; Strome, B.; Ba, A. N. N.; Alberti, S.; Forman-Kay, J. D.; Moses, A. M. Proteome-Wide Signatures of Function in Highly Diverged Intrinsically Disordered Regions. *Elife* **2019**, *8*, e46883. https://doi.org/10.7554/elife.46883.

(35) Hunter, J. D. Matplotlib: A 2D Graphics Environment. *Comput. Sci. Eng.* **2007**, *9* (3), 90–95. https://doi.org/10.1109/mcse.2007.55.

(36) *Protein Calculator v3.4*. https://protcalc.sourceforge.net/ (accessed 2025-06-24).

(37) Kabsch, W.; Sander, C. Dictionary of Protein Secondary Structure: Pattern Recognition of Hydrogen-bonded and Geometrical Features. *Biopolymers* **1983**, *22* (12), 2577–2637. https://doi.org/10.1002/bip.360221211.
